# Spatiotemporal distribution of ROS production, delivery and utilization in Arabidopsis root hairs

**DOI:** 10.1101/2023.01.05.522872

**Authors:** Lenka Kuběnová, Jan Haberland, Petr Dvořák, Jozef Šamaj, Miroslav Ovečka

## Abstract

Fluorescent selective probes for reactive oxygen species (ROS) detection in living cells are versatile tools for the documentation of ROS production in plant developmental or stress reactions. We employed high-resolution live-cell imaging and semi-quantitative analysis of *Arabidopsis thaliana* stained with CM-H_2_DCFDA, CellROX^TM^ Deep Red and Amplex^TM^ Red for functional characterization of spatiotemporal mode of ROS production, delivery and utilization during root hair formation. Cell viability marker fluorescein diacetate served as a positive control for dye-loading and undisturbed tip growth after staining. Colocalization analysis with subcellular molecular markers and utilization of two root hair mutants with similar phenotype of non-elongating root hairs, but with contrast reasons for this impairment, we found that: i) CM-H_2_DCFDA is a sensitive probe for ROS generation in cytoplasm, ii) CellROX^TM^ Deep Red labels ROS in mitochondria, iii) Amplex^TM^ Red labels apoplastic ROS and mitochondria, and shows high selectivity to root hairs, iv) *rhd2-1* mutant with nonfunctional AtRBOHC/RHD2 has a low level of CM-H_2_DCFDA-reactive ROS in cytoplasm and lacks Amplex^TM^ Red-reactive ROS in apoplast, v) *ACTIN2*-deficient *der1-3* mutant is not altered in these aspects. The sensitivity of CellROX^TM^ Deep Red was documented by discrimination between larger ROS-containing mitochondria and small, yet ROS-free pre-mature mitochondria in the growing tip of root hairs. We characterized spatial changes in ROS production and compartmentalization induced by external ROS modulators, ethylene precursor 1-aminocyclopropane-1-carboxylic acid and ionophore valinomycin. This dynamic and high-resolution study of ROS production and utilization opens new opportunities for precise speciation of particular ROS involved in the root hair formation.

**One sentence summary:** High-resolution live-cell imaging of ROS production and subcellular localization in bulges and growing root hairs of Arabidopsis using CM-H_2_DCFDA, CellROX^TM^ Deep Red and Amplex^TM^ Red selective probes.

## Introduction

Root hairs are tubular outgrowths of specialized root epidermal cells, trichoblasts. A specific feature of the formation of root hairs is a spatiotemporal change in the polarity, driving cell expansion. From the trichoblast expanding by diffuse cell growth, a fundamental change in expansion mode occurs, and the root hair itself is further elongated only by polarized tip growth (Dolan et al., 1994; Baluška et al., 2000), a special mode of extension also shared by pollen tubes (Schoenaers et al., 2017). Root hairs increase the root’s total surface, significantly improving water and nutrient uptake from soil. They also support the anchoring of the plant in the soil and enable symbiotic interactions with the beneficial microflora in the rhizosphere (Gilroy and Jones, 2000). The establishment and regulation of the tip growth is conditioned by the maintenance of the polar organization of the tip-growing cell. Membrane transport processes, temporal and spatial regulation of exocytosis and endocytosis, as well as selective recycling of vesicles play a central role in this maintenance. Polar tip growth, including the maintenance of root hair polarity itself, requires regulated directional transport, dynamic properties of the cytoskeleton, maintenance of physiological gradients in the tip and, last but not least, signaling cascades (Baluška et al., 2000; Šamaj et al., 2006). Pollen tubes and root hairs are model objects for the study of tip growth in plants (Schoenaers et al., 2017).

The formation of a bulge on the outer cell wall of the trichoblast represents a transitional phase between the diffuse growth of the epidermal cell and the tip growth of the root hair. From a mechanical and functional point of view, it is a rebuilding of the cell wall, which occurs primarily by local loosening of bonds. There is a local decrease in the pH of the cell wall (Bibikova et al., 1998), and a localized increase in the concentration of protons in the cytoplasm (Bibikova et al., 1997). This is associated with increased xyloglucan endotransglycosylase activity (Vissenberg et al., 2001) and specific expression of expansin 7 and 10 in trichoblasts (Cho and Cosgrove, 2002). The creation of a proton gradient is also closely connected with the production of reactive oxygen species (ROS; Foreman et al., 2003). The tip growth mechanism of root hairs requires a tip-focused gradient of Ca^2+^ in the cytosol, which regulates the activity of vesicular trafficking, generation of ROS, and organization of the cytoskeleton in the growing tip. Ca^2+^-permeable ion channels are localized at the apical plasma membrane (Miedema et al., 2008) and ROS are generated by *A. thaliana* respiratory burst oxidase homolog protein C/ ROOT HAIR DEFECTIVE 2 (AtrbohC/RHD2). The ROS production is regulated by ROP GTPases (Foreman et al., 2003) and the whole process is interconnected by a positive feedback mechanism, which determines root hair polarity and tip growth (Carol et al., 2005; Takeda et al., 2008). Tip-focused accumulation of ROS is also tightly associated with pollen tube tip growth (Schoenaers et al., 2017). AtABCG28, an ABC transporter, is responsible for ROS accumulation at the tip of the growing *Arabidopsis thaliana* pollen tubes (Do et al., 2019). In tip-growing cells, NADPH oxidases, highly regulated transmembrane proteins that use cytosolic NADPH, produce superoxide (O_2_^•−^) at the apoplastic side of the plasma membrane, which is then converted to hydrogen peroxide (H_2_O_2_) by the activity of the superoxide dismutase (SOD) in the apoplast (Demidchik, 2018). Hydrogen peroxide can diffuse to the cytoplasm of the tip-growing cells via aquaporins, forming a tip-focused gradient of ROS (Bienert and Chaumont, 2014).

For quantitation of H_2_O_2_ in living cells quite extensively is used the oxidation of 2’-7’ dichlorofluorescin (H_2_DCF) to 2’-7’dichlorofluorescein (DCF). An important parameter is the good permeability of the diacetate form, H_2_DCFDA and its acetomethyl ester H_2_DCFDA-AM through plasma membrane of living cells. Probes are taken up by cells and nonspecific cellular esterases cleave off the lipophilic groups, resulting in trapping of the charged compound inside the cell. Oxidation of H_2_DCF by ROS in cells converts the molecule to 2’,7’ dichlorodihydrofluorescein (DCF), which is highly fluorescent. Although DCF was believed to be specific for H_2_O_2_, some evidence indicates that other radicals, such as peroxynitrite and hypochlorous acid can oxidize H_2_DCF (Hoffman et al., 2008). However, albeit this probe can react with other ROS, due to its high sensitivity, it has been widely used to monitor H_2_O_2_ production in plants (Swanson et al., 2011). The CellROX^TM^ Deep Red is a fluorescent reagent that detects O_2_ ^•−^ and hydroxyl radicals (Alves et al., 2015). It is a cell-permeant dye weakly fluorescent while in a reduced state. However, it exhibits strong and photostable fluorescence upon oxidation by ROS. Amplex^TM^ Red is a most commonly used fluorogenic substrate, which serves as hydrogen donors in conjugation with horseradish peroxidase (HRP) enzyme to produce intensely fluorescent product. Amplex^TM^ Red serves a potent and sensitive probe in this case, as increasing amounts of H_2_O_2_ in cells leads to increasing amounts of specific fluorescent product. Amplex^TM^ Red is oxidized by H_2_O_2_ in the presence of HRP and converts to resorufin (Reszka et al., 2005).

In tip-growing cells, fluorescence localization of ROS using CM-H_2_DCFDA or DCFH_2_-DA was quite commonly employed. In pollen tubes, it was utilized for H_2_O_2_ detection (Potocký et al., 2007; Do et al., 2019). The same approach was used for the localization of ROS production in Arabidopsis roots upon salt stress-induced signal transduction (Leshem et al., 2007) and in root hairs of Arabidopsis upon treatment with LY294002, a phosphatidylinositol 3-kinase (PI3K) - specific inhibitor (Lee et al., 2008). Alternatively, to test the possibility that the ROS produced by mitochondria might influence the tip growth of pollen tubes, a CellROX^TM^ Deep Red probe, detecting O_2_^•−^ and hydroxyl radicals was used (Do et al., 2019). In the absence of ROS, it remains in its reduced state with no fluorescence, which is rapidly induced upon oxidation during cellular oxidative stress. It has been shown that CellROX^TM^ Deep Red can be used to detect ROS production in ram sperm by *in vitro* and *in vivo* oxidative stress induction (Alves et al., 2015). To our knowledge, Amplex^TM^ Red was not used to study ROS production, distribution and utilization during the root hair formation.

Sensitive fluorescent probes were so far not utilized for subcellular discrimination between ROS sources in growing root hairs. Mapping of the origin, subcellular location of the production and dynamics of ROS utilization during bulge formation and tip growth of root hairs has not been documented and remains unknown. In this study, live-cell imaging using high-resolution spinning disk fluorescence microscopy supplemented with semi-quantitative analysis was used for functional characterization of spatiotemporal mode of ROS production, delivery and utilization in Arabidopsis root hairs from bulge formation to tip growth. Staining of *Arabidopsis thaliana* root apex with probes for selective ROS detection in living cells, CM-H_2_DCFDA, CellROX^TM^ Deep Red and Amplex^TM^ Red, showed different subcellular staining patterns. Since the first screening analysis revealed that staining patterns of these three ROS-selective probes in growing root hairs may differ, we analyzed subcellular ROS production also in *rhd2-1* and *der1-3* root hair mutants. To prove the sensitivity of the used probes, we also determined spatial changes in ROS production and compartmentalization in root hairs induced by external ROS modulators such as ACC and valinomycin. Overall, the approach of detailed subcellular qualitative and quantitative ROS distribution patterns by using sensitive probes in living cells may serve as a new platform for a deeper understanding of root hair development.

## Results

### Fluorescence ROS detection in Arabidopsis root hairs by live-cell imaging

Microscopic visualization of growing root hairs, their non-destructive loading with fluorescent probes, and subsequent fluorescence live-cell imaging for tens of minutes is possible only at physiologically-relevant conditions. This is fundamental not only when fluorescently-detected ROS will be visualized and measured, but also in control media before the probe loading. In our system, together with checking a normal root hair morphology and typical intracellular polarity, a continuation of tip growth has been monitored in all experiments. To this point, as a positive control for fluorescent dyes loading and performance of treated plants in the microscope, staining with a cell viability marker, fluorescein diacetate (FDA) was used. The probe labeled all cells in the root tip and root hair formation zone of Col-0 (Figure 1A), *rhd2-1* (Figure 1B) and *der1-3* (Figure 1C) roots. A prominent and strong signal was detected in growing root hairs of Col-0 plants, accumulated mainly in apical and subapical zones of root hairs (Figure 1A). Detailed analysis revealed labeling of the cytoplasm in growing root hairs of Col-0 (Figure 1D) and C24 (Figure 1E) wild-types, supplemented by a signal located in distinct subcellular spots. Root hairs of *rhd2-1* and *der1-3* mutants cannot elongate by a tip growth (Figure 1B,C). Therefore, only bulges were present, showing typical fluorescence in the cytoplasm after FDA staining in *rhd2-1* (Figure 1F) and *der1-3* (Figure 1G) roots. Analysis of root hair FDA fluorescence at time points of 0 min (Figure 1H,K), 5 min (Figure 1I,L) and 10 min (Figure 1J,M) of imaging revealed equal distribution of the fluorescence signal in apical and subapical parts of growing root hairs with negligible temporal variations (Figure 1N). This favors FDA serving as a positive control for dye-loading, showing that any difference in signal intensity between time-points of 0, 5 and 10 minutes, or mutants observed in further experiments are not related to prolonged or unequal dye accessibility or internalization during imaging. Root hair tip growth of untreated plants observed in time-lapsed imaging mode using 100× lens in the spinning disk microscope was not disturbed (Supplemental Movie S1). FDA staining partially reduced the root hair tip growth rate as compared to unstained Col-0 root hairs, but values around 1 µm·min^-1^ recorded after FDA staining (Figure 1O) lay within a typical range of Arabidopsis root hair growth rate.

**Figure 1.**
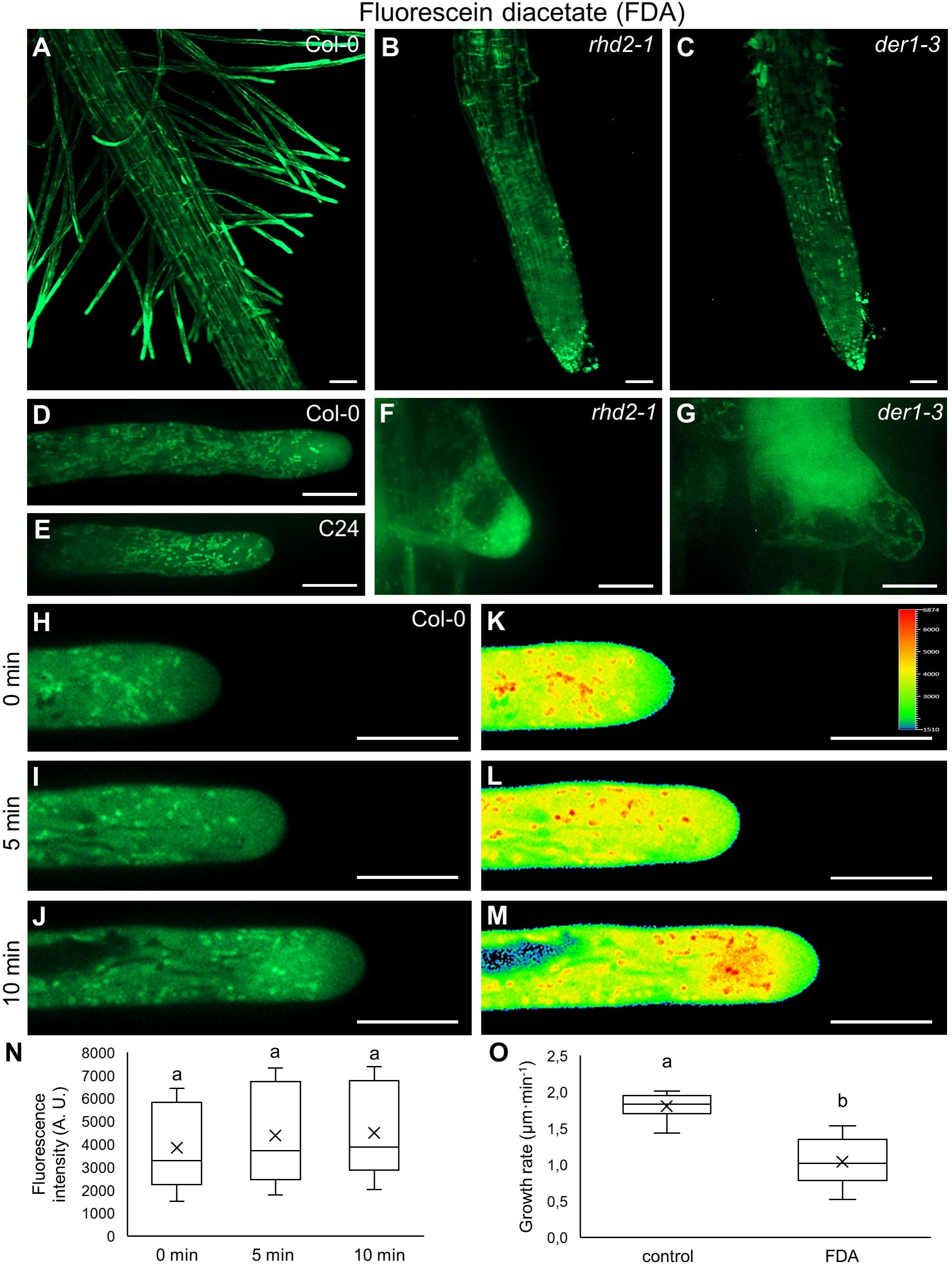
Monitoring of root hairs, their viability and tip growth during imaging by staining with FDA. **A-C.** Bulges and growing root hairs in root hair formation zone of Col-0 **(A)**, *rhd2-1* **(B)** and *der1-3* **(C)** roots. **(D-G).** FDA staining of growing root hairs of Col-0 **(D)**, C24 **(E)** and bulges of *rhd2-1* **(F)** and *der1-3* **(G)** mutants. **H-M.** Growing root hair of Col-0 after FDA staining **(H­ J)**, and pseudo-color-coded fluorescence intensity visualization **(K-M).** Root hair was imaged at time points of O min **(H,K)**, 5 min **(l,L)** and 10 min **(J,M)** of growth. Fluorescence intensity distribution is visualized in a pseudo-color-coded scale, where dark blue represents minimal intensity and red represents maximum intensity (inset in **K). N.** Mean fluorescence intensity of FDA measured in root hair tips (measured area schematically illustrated in Supplemental Figure S1B) at imaging times of O min, 5 min, and 10 min. **0.** Averaged root hair tip growth rate of control and FDA-treated root hairs of Col-0 plants measured during imaging within the time period of 10 min. N = 9-10. Box plots display the first and third quartiles, split by the median; the crosses indicate the mean values; whiskers extend to include the max/min values. Lowercase letters indicate statistical significance between lines according to one­ way ANOVA with Fisher’s LSD tests (P < 0.05). Scale bar= 50 µm **(A-C)**, 10 µm **(D-M).**

We analyzed fluorescence signal distribution in apical parts of developing bulges and growing root hairs after ROS staining with CM-H_2_DCFDA, CellROX^TM^ Deep Red and Amplex^TM^ Red probes by qualitative and semi-quantitative analyses. We characterized a fluorescence intensity distribution measured along a line profile oriented longitudinally at the apex of bulges and root hairs that was 10 µm long and reached the apical cell wall (Supplemental Figure S1A). Mean fluorescence intensity was measured in a 10 µm segment encompassing a clear zone and sub-apical part of bulges and root hairs (Supplemental Figure S1B). Mean fluorescence signal intensity in the area covering only the cell wall was measured within the clear zone and sub-apical part of root hairs (Supplemental Figure S1C). Furthermore, mean fluorescence signal intensity in the zone of small organelles distribution was measured in the area located in the subapical region of root hairs (Supplemental Figure S1D). Within the 10 min-long time-lapsed imaging with images captured every 30 s, individual frames from three imaging time points (0 min, 5 min and 10 min) are presented in each experiment.

### Differences in ROS detection in bulges and root hairs of control plants and root hair mutants

Staining with CM-H_2_DCFDA showed that all epidermal cells in the elongation and differentiation zones of Col-0 roots were stained, while prominent labeling appeared in bulges, short root hairs and apices of longer growing root hairs (Supplemental Figure S2A-B). In developing bulges, staining with CM-H_2_DCFDA revealed the presence of a relatively low signal, distributed evenly in the cytoplasm without any preference to particular subcellular compartments (Supplemental Figure S2E-G; Supplemental Movie S2). An overview of the root hair formation zone revealed a prominent labeling pattern observed in apices of growing root hairs (Supplemental Figure S2A). Therefore, after documentation of signal intensity and distribution in bulges, we analyzed the subcellular distribution of fluorescence signal in growing root hairs. Staining with CM-H_2_DCFDA led to the accumulation of specific signal in cytoplasm, which was particularly prominent in root hair apical and subapical zones and decreased in the apical clear zone (Figure 2A-C; Supplemental Figure S2H-J). Analysis at time points 0 min, 5 min and 10 min from the beginning of imaging showed that in growing root hairs the zone with the highest signal was still located in their apical and subapical parts (Figure 2A-C; Supplemental Figure S2H-J; Supplemental Movie S3).

**Figure 2.**
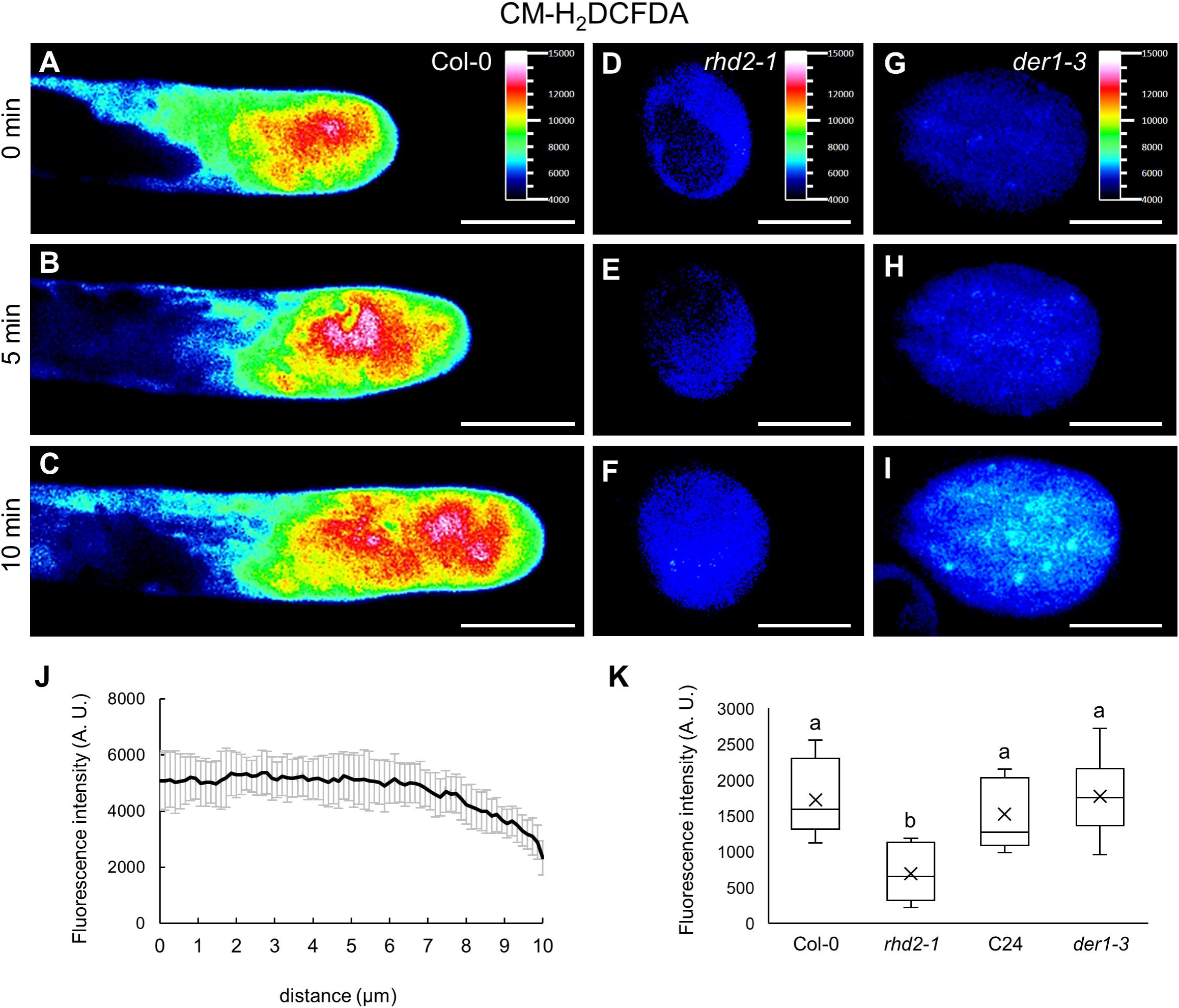
Distribution of ROS in root hairs of Col-0 and bulges of *rhd2-1* and *der1-3* mutants after staining with CM-H_2_DCFDA. **A-I.** Growing root hair of Col-0 **(A-C)**, developing bulge of *rhd2-1* mutant **(D-F)** and *der1-3* mutant **(G-1).** Root hairs and bulges were imaged at time points of O min **(A,D,G)**, 5 min **(B,E,H)** and 10 min **(C,F,I)** of growth. Fluorescence intensity distribution is visualized in a pseudo-color-coded scale, where black represents minimal intensity and white represents maximum intensity (insets in **A,D,G). J.** Mean fluorescence intensity distribution of CM-H_2_DCFDA measured along a 10 µm line oriented longitudinally at the central part of Col-0 growing root hairs, reaching the apical cell wall (schematically illustrated in Supplemental Figure S1A). **K.** Semi-quantitative mean CM-H_2_DCFDA fluorescence intensity analysis (measured area schematically illustrated in Supplemental Figure S1B) in bulges of *rhd2-1* and *der1-3* mutants, and respective Col-0 and C24 wild-types. N = 10-21. Box plots display the first and third quartiles, split by the median; the crosses indicate the mean values; whiskers extend to include the max/min values. Lowercase letters indicate statistical significance between lines according to one-way ANOVA with Fisher’s LSD tests (P < 0.05). Scale bar= 10 µm **(A-1).**

To reveal the relationship between subcellular sources of ROS generation and the mechanism of root hair tip growth, we analyzed two root hair mutants with phenotype of short root hairs (bulges) that cannot elongate by tip growth. First *rhd2-1* mutant (*root hair defective 2-1*) bears a loss-of-function mutation in the *AtRBOHC/RHD2* locus, resulting in missing tip-focused ROS and Ca^2+^ gradients (Schiefelbein and Somerville, 1990; Foreman et al., 2003). Second *der1-3* (*deformed root hairs 1-3*) mutant bears a single-point mutation in the *ACTIN2* gene and shows impairment in root hair elongation after bulge establishment (Ringli et al., 2002). Fluorescence ROS staining with CM-H_2_DCFDA revealed a very low signal in the root hair formation zone of *rhd2-1* mutant (Supplemental Figure S2C), while it was prominent in this zone of *der1-3* mutant (Supplemental Figure S2D). Live-cell imaging of ROS in bulges of *rhd2-1* mutant after staining with CM-H_2_DCFDA revealed a very low signal in the cytoplasm that remained unchanged in recording time points at 0 min, 5 min and 10 min of imaging (Figure 2D-F; Supplemental Figure S2K-M; Supplemental Movie S4). Unlikely, the fluorescence signal after staining with CM-H_2_DCFDA was considerably stronger in bulges of *der1-3* mutant (Figure 2G-I; Supplemental Figure S2N-P; Supplemental Movie S5). Analysis of mean fluorescence intensity distribution of CM-H_2_DCFDA along a profile (Supplemental Figure S1A) in growing Col-0 root hairs showed equal distribution of the fluorescence signal in apical and subapical zones, with decreasing tendency in the clear zone (Figure 2J). Accordingly, mean fluorescence intensity measurement (Supplemental Figure S1B) in bulges of *rhd2-1* and *der1-3* mutants and their respective Col-0 and C24 wild-types revealed a significantly reduced amount in *rhd2-1*, while CM-H_2_DCFDA fluorescence in bulges of Col-0, C24 and *der1-3* was higher and similar to each other (Figure 2K). Fluorescence intensity measurement along the profile in apical parts of bulges and root hairs after ROS staining with CM-H_2_DCFDA in Col-0 plants was acquired during their growth (Supplemental Figure S3A; Supplemental Movie S6, S7). Distribution of fluorescence signal after line profile measurement in growing bulges (Supplemental Figure S3B), short root hairs reaching the range of 10-200 µm (Supplemental Figure S3C), and long root hairs reaching the length over 200 µm (Supplemental Figure S3D) did not show differences among measured 0 min, 5 min and 10 min time-points of imaging. However, profile fluorescence intensities increased during root hair development from bulges (Supplemental Figure S2E-G) to long root hairs (Supplemental Figure S2H-J, S3E). This analysis also revealed that CM-H_2_DCFDA-specific signal was distributed equally in the subapical region, decreasing considerably in the apical clear zone. Semi-quantitative mean CM-H_2_DCFDA fluorescence intensity analysis in growing root hairs of Col-0 and C24 wild-types did not show considerable differences (Supplemental Figure S3F).

Staining with CellROX^TM^ Deep Red led to the detection of fluorescence signal in root epidermal cells of Col-0 plants, and it increased in bulges and apices of growing root hairs (Supplemental Figure S4A-B). Comparable fluorescent staining with CellROX^TM^ Deep Red was observed also in root hair formation zone of *rhd2-1* (Supplemental Figure S4C) and *der1-3* (Supplemental Figure S4D) mutants. After staining with CellROX^TM^ Deep Red, resulting signal was localized in distinct intracellular oval compartments in developing bulges. The size, number and cellular distribution of ROS-positive compartments were not changed during bulge development, and this staining pattern was stable over recorded time points (Supplemental Figure S4E-G; Supplemental Movie S8). In growing root hairs stained with CellROX^TM^ Deep Red these compartments were distributed in the cytoplasm within the subapical zone and around the vacuole, but were absent in clear zone (Figure 3A-C; Supplemental Figure S4H-J). The most prominent accumulation of these compartments was visible in root hair subapical zone (Figure 3A-C; Supplemental Figure S4H-J; Supplemental Movie S9). Staining of ROS with CellROX^TM^ Deep Red in root hair mutants showed a similar pattern as observed in bulges of wild-type. CellROX^TM^ Deep Red-dependent ROS localization in compartments moving in the cytoplasm was observed in bulges of both *rhd2-1* (Figure 3D-F; Supplemental Figure S4K-M; Supplemental Movie S10) and *der1-3* (Figure 3G-I; Supplemental Figure S4N-P; Supplemental Movie S11) mutants.

**Figure 3.**
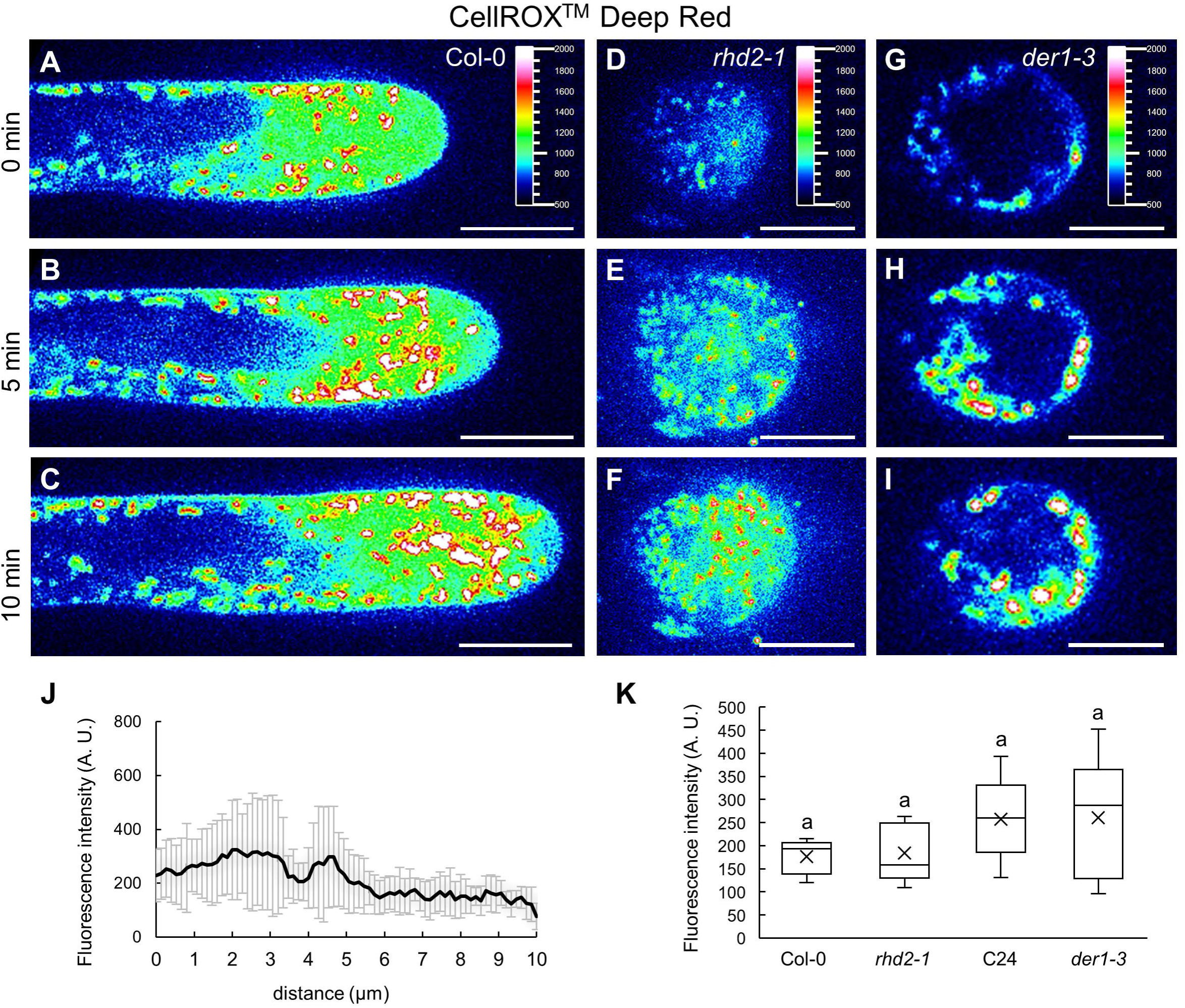
Distribution of ROS in root hairs of Col-0 and bulges of *rhd2-1* and *der1-3* mutants after staining with CellROX™ Deep Red. **A-I.** Growing root hair of Col-0 **(A-C)**, developing bulge of *rhd2-1* mutant **(D-F)** and *der1-3* mutant **(G-1).** Root hairs and bulges were imaged at time points of O min **(A,D,G)**, 5 min **(B,E,H)** and 10 min **(C,F,I)** of growth. Fluorescence intensity distribution is visualized in a pseudo-color­ coded scale, where black represents minimal intensity and white represents maximum intensity (insets in **A,D,G). J.** Mean fluorescence intensity distribution of CellROX™ Deep Red measured along a 10 µm line oriented longitudinally at the central part of Col-0 growing root hairs, reaching the apical cell wall (schematically illustrated in Supplemental Figure S1A). **K.** Semi-quantitative mean CellROX™ Deep Red fluorescence intensity analysis (measured area schematically illustrated in Supplemental Figure S1B) in bulges of *rhd2-1* and *der1-3* mutants, and respective Col-0 and C24 wild-types. N = 10-17. Box plots display the first and third quartiles, split by the median; the crosses indicate the mean values; whiskers extend to include the max/min values. Lowercase letters indicate statistical significance between lines according to one­ way ANOVA with Fisher’s LSD tests (P < 0.05). Scale bar= 10 µm **(A-1).**

Mean fluorescence intensity distribution in CellROX^TM^ Deep Red-stained growing root hairs of Col-0 plants showed fluctuation of the fluorescence signal intensity, which was caused by the distribution pattern of ROS-positive subcellular compartments that were actively moving. It was visible mainly in the subapical zones, but not in the clear zones of root hairs (Figure 3J; Supplemental Figure S5A). A similar pattern of distribution and active movement of CellROX^TM^ Deep Red-stained subcellular compartments was recorded in bulges and the pattern was stable among recorded time points at 0 min, 5 min and 10 min (Supplemental Figure S5B; Supplemental Movie S12). Compared to bulges, the mean fluorescence intensities considerably increased in growing root hairs. Due to the active movement of ROS-positive compartments, fluctuations in their mean fluorescence intensities occurred and were obvious among recorded time points at 0 min, 5 min and 10 min (Supplemental Figure S5C-E; Supplemental Movie S13). Similarly to the staining pattern by CM-H_2_DCFDA, the signal intensity after CellROX^TM^ Deep Red staining decreased in the apical part of root hairs and was considerably low in their clear zones (Figure 3J; Supplemental Figure S5C-E). Mean fluorescence intensity measured in the whole apical part of bulges in *rhd2-1* and *der1-3* mutants, as well as in their respective Col-0 and C24 wild-types, revealed no differences (Figure 3K). In growing root hairs of Col-0 and C24, the mean fluorescence intensity of CellROX^TM^ Deep Red staining in whole apical and subapical parts was higher in C24 genotype (Supplemental Figure S5F).

Staining of Col-0 roots with Amplex^TM^ Red revealed that root epidermal cells were virtually negative, unlike CM-H_2_DCFDA and CellROX^TM^ Deep Red probes. However, bulges, short root hairs, and apices of longer and still growing root hairs were specifically labeled by Amplex^TM^ Red (Supplemental Figure S6A-B). The fluorescence signal of ROS stained with Amplex^TM^ Red was very low in root hair formation zone of *rhd2-1* mutant (Supplemental Figure S6C). On the other hand, it was strong and comparable to Col-0 in root hair formation zone of *der1-3* mutant (Supplemental Figure S6D). Imaging of the root hair formation zone in Col-0 at higher resolution revealed that developing bulges were outlined by Amplex^TM^ Red-specific signal at the surface (Supplemental Figure S6E-G; Supplemental Movie S14). In growing root hairs, the Amplex^TM^ Red signal also outlined root hairs at the surface, together with subcellular compartments showing weak fluorescence and distributed in the cytoplasm within the subapical part of root hairs (Figure 4A-C; Supplemental Figure S6H-J; Supplemental Movie S15). Staining of root hair bulges with Amplex^TM^ Red provided almost negative results in *rhd2-1* mutant (Figure 4D-F; Supplemental Figure S6K-M; Supplemental Movie S16), while prominent fluorescent signal outlined the surface of bulges in *der1-3* mutant (Figure 4G-I; Supplemental Figure S6N-P; Supplemental Movie S17).

**Figure 4.**
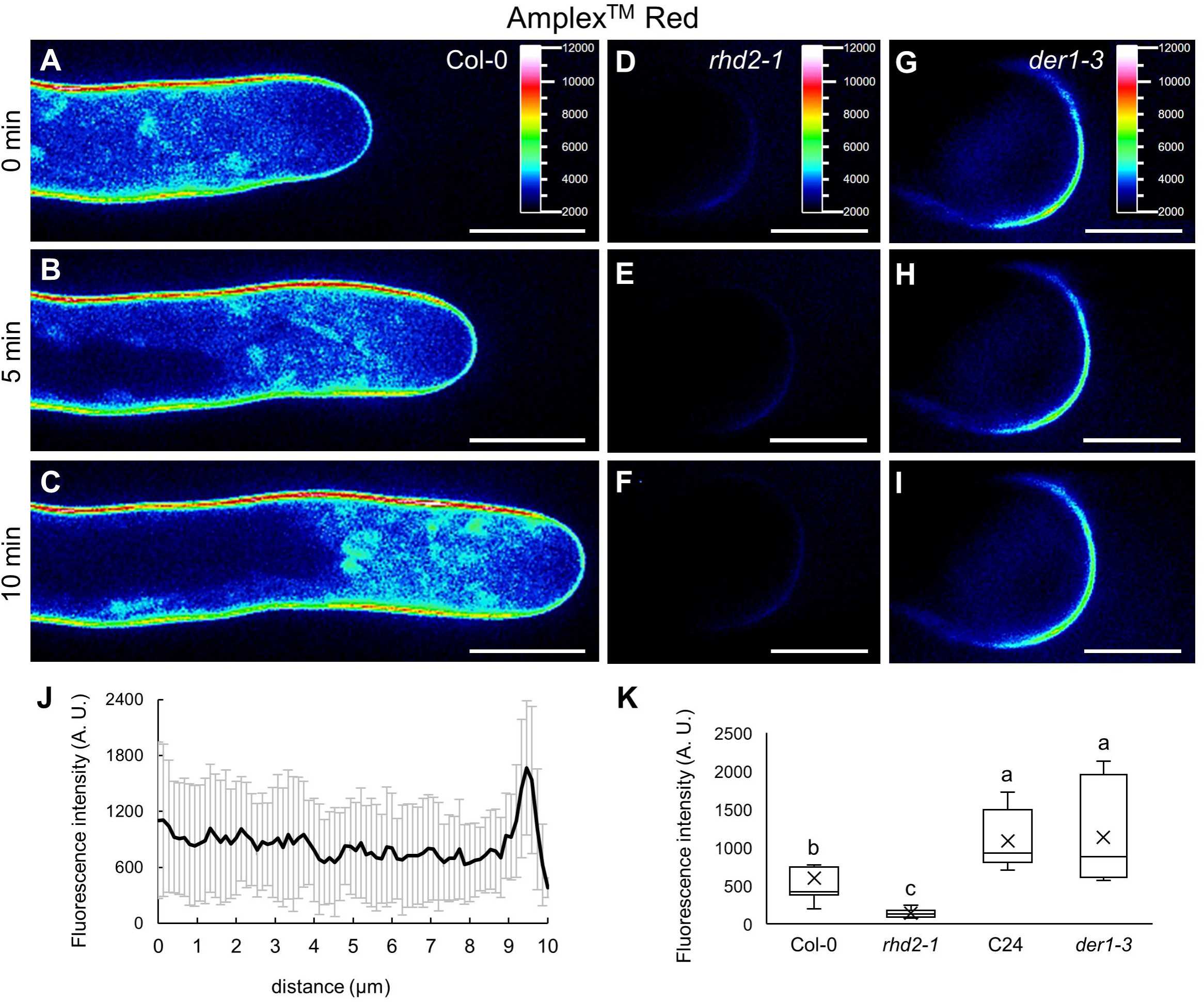
Distribution of ROS in root hairs of Col-0 and bulges of *rhd2-1* and *der1-3* mutants after staining with Amplex™ Red. **A-I.** Growing root hair of Col-0 **(A-C)**, developing bulge of *rhd2-1* mutant **(D-F)** and *der1-3* mutant **(G-1).** Root hairs and bulges were imaged at time points of O min **(A,D,G)**, 5 min **(B,E,H)** and 10 min **(C,F,I)** of growth. Fluorescence intensity distribution is visualized in a pseudo-color-coded scale, where black represents minimal intensity and white represents maximum intensity (insets in **A,D,G). J.** Mean fluorescence intensity distribution of Amplex^TM^ Red measured along a 10 µm line oriented longitudinally at the central part of Col-0 growing root hairs, reaching the apical cell wall (schematically illustrated in Supplemental Figure S1A). **K.** Semi-quantitative mean Amplex™ Red fluorescence intensity analysis (measured area schematically illustrated in Supplemental Figure S1C) in bulges of *rhd2-1* and *der1-3* mutants, and respective Col-0 and C24 wild-types. N = 10. Box plots display the first and third quartiles, split by the median; the crosses indicate the mean values; whiskers extend to include the max/min values. Lowercase letters indicate statistical significance between lines according to one-way ANOVA with Fisher’s LSD tests (P < 0.05). Scale bar= 10 µm **(A-1).**

Mean fluorescence intensity distribution within the apex of growing Col-0 root hairs after staining with Amplex^TM^ Red, analyzed in the distance of 10 µm longitudinally from the apical cell wall (Supplemental Figure S7A), revealed a prominent signal located at the root hair periphery in the tip, indicating the cell wall (Figure 4J). A similar pattern of fluorescence signal distribution at the surface was recorded in bulges (Supplemental Figure S7B; Supplemental Movie S18) and growing (both short and long) root hairs (Supplemental Figure S7C-D; Supplemental Movie S19) with no differences among recorded time points at 0 min, 5 min and 10 min. The signal intensity, however, increased from bulges to growing root hairs (Supplemental Figure S7E). Mean fluorescence intensity measured in the cell wall of apical bulge parts (the area of measurement schematically illustrated in Supplemental Figure S1C) showed very low level in *rhd2-1* mutant (Figure 4K). It was significantly higher in *der1-3* mutant, which was along with C24 also higher in comparison to Col-0 (Figure 4K). This difference was corroborated also by comparison of Col-0 and C24 alone, showing higher mean area fluorescence intensity in C24 (Supplemental Figure S7F).

### Subcellular identity of ROS-positive compartments in root hairs labeled with CellROX^TM^ Deep Red

In developing bulges and growing root hairs, staining with CM-H_2_DCFDA revealed, that the fluorescence signal distributed evenly in cytoplasm without any preference to particular subcellular compartments (Figure 2A-C; Supplemental Figure S2H-J; Supplemental Movie S3). However, CellROX^TM^ Deep Red stained motile compartments distributed in cytoplasm of root hairs with the exception of the clear zone (Supplemental Figure S4H-J; Supplemental Movie S9). This prompted us to define their nature by using colocalization analysis with known subcellular molecular markers. We have used transgenic lines carrying i) GFP-RHD2, a GFP-tagged AtRBOHC/RHD2 (Takeda et al., 2008; Kuběnová et al., 2022), ii) GFP-RabA1d marker for early endosomal/TGN compartments (Ovečka et al., 2010; Berson et al., 2014; von Wangenheim et al., 2016), iii) RabF2a-YFP marker of late endosomes (von Wangenheim et al., 2016), and iv) GFP-tagged mitochondria-targeting sequence of the N-terminus of the F1-ATPase g-subunit (Niwa et al., 1999). GFP-RHD2 locates to vesicular compartments belonging to early endosomes/TGN and to apical plasma membrane in growing root hairs (Figure 5A). The detailed view did not reveal a colocalization between ROS-positive compartments and GFP-RHD2 (Figure 5B), which was confirmed by a fluorescence intensity profile measurement through the compartments (Figure 5C). GFP-RabA1d was located in early endosomal/TGN compartments accumulated in the apical and subapical regions of root hairs (Figure 5D) and detailed analysis showed no colocalization with ROS-positive compartments (Figure 5E-F). RabF2a-YFP decorated larger late endosomes located mainly away from the apical and subapical regions of root hairs (Figure 5G). Higher magnification (Figure 5H) and a semi-quantitative profile measurement of fluorescence signal intensities (Figure 5I) revealed no colocalization with ROS-positive compartments. On the other hand, mitochondria identified by a mitochondrial GFP marker showed the same size, shape and distribution pattern as ROS-positive compartments after CellROX^TM^ Deep Red staining (Figure 5J). A high degree of their colocalization was revealed by both high magnification visualization (Figure 5K) and fluorescence intensity profile measurements (Figure 5L).

**Figure 5.**
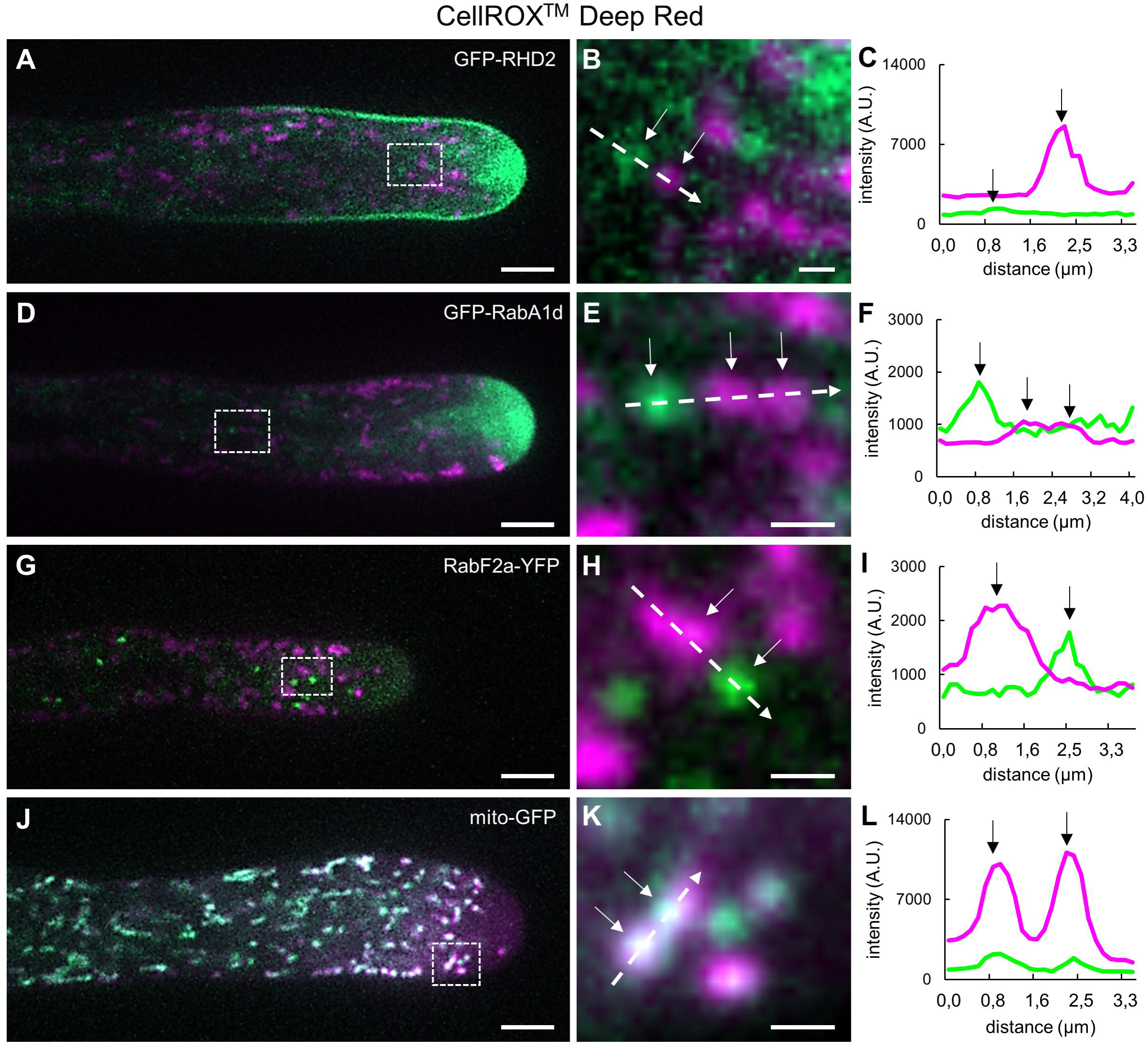
Colocalization analysis of subcellular molecular markers in root hairs with compartments accumulating ROS after staining with CellROX™ Deep Red. **A-C.** Colocalization with transgenic line for visualization of vesicular compartments containing GFP-RHD2, a GFP-tagged AtRBOHC/RHD2. **D-F.** Colocalization with transgenic line carrying early endosomal marker GFP-RabA1d. **G-1.** Colocalization with transgenic line carrying late endosomal marker RabF2a-YFP. **J-L.** Colocalization with transgenic line carrying GFP-tagged mitochondria. Overview of root hairs carrying molecular markers (in green) and ROS­ accumulating compartments (in magenta) after CellROX™ Deep Red staining **(A,D,G,J).** Detailed images of compartments containing carrying molecular markers and ROS **(B,E,H,K)** from the white boxes shown in **(A,D,G,J).** Fluorescence intensity profiles **(C,F,l,L)** of fluorescence signals of molecular markers (green lines) and ROS-containing compartments (magenta lines) along the interrupted white lines shown in **(B,E,H,K).** Arrows in **(B,C,E,F,H,l,K,L)** indicate the position of the compartments. Scale bar = 5 µm **(A,D,G,J)**, 1 µm **(B,E,H,K).**

High-resolution time-lapse video-recordings of growing root hairs in Arabidopsis plants expressing a mitochondrial GFP marker and stained with CellROX^TM^ Deep Red at a given frequency of image acquisition every 30 sec revealed dynamic pattern of colocalization (Figure 6A-C). We used a single particle dynamic colocalization analysis to track events when signal of ROS accumulation after staining with CellROX^TM^ Deep Red colocalized with mitochondria. However, some GFP-labelled mitochondria appeared also temporarily without CellROX^TM^ Deep Red-based ROS staining. This occurred mainly in the clear zone and just behind the clear zone of growing root hairs, but not deeper in the subapical and shank regions of root hairs (Figure 6A-C). In general, the clear zone of growing root hairs was free of mitochondria and only sparsely they entered this region. Such GFP-labeled but initially ROS-free mitochondria were small, round and highly motile. Therefore, they temporarily showed no CellROX^TM^ Deep Red signal at the beginning, which was replenished after a short time (Figure 6A,D). Also events resembling small round-shaped mitochondria separation from larger elongated ones already containing ROS stained with CellROX^TM^ Deep Red (“fission”) were observed. In such cases, small newly-separated mitochondria were not stained with CellROX^TM^ Deep Red, however, they acquired it shortly after appearance (Figure 6B,E). Generally, ROS-free appearance was typical for small mitochondria showing dynamic movement that were visually different from larger and less mobile mitochondria already containing ROS stained with CellROX^TM^ Deep Red (Figure 6C,F).

**Figure 6.**
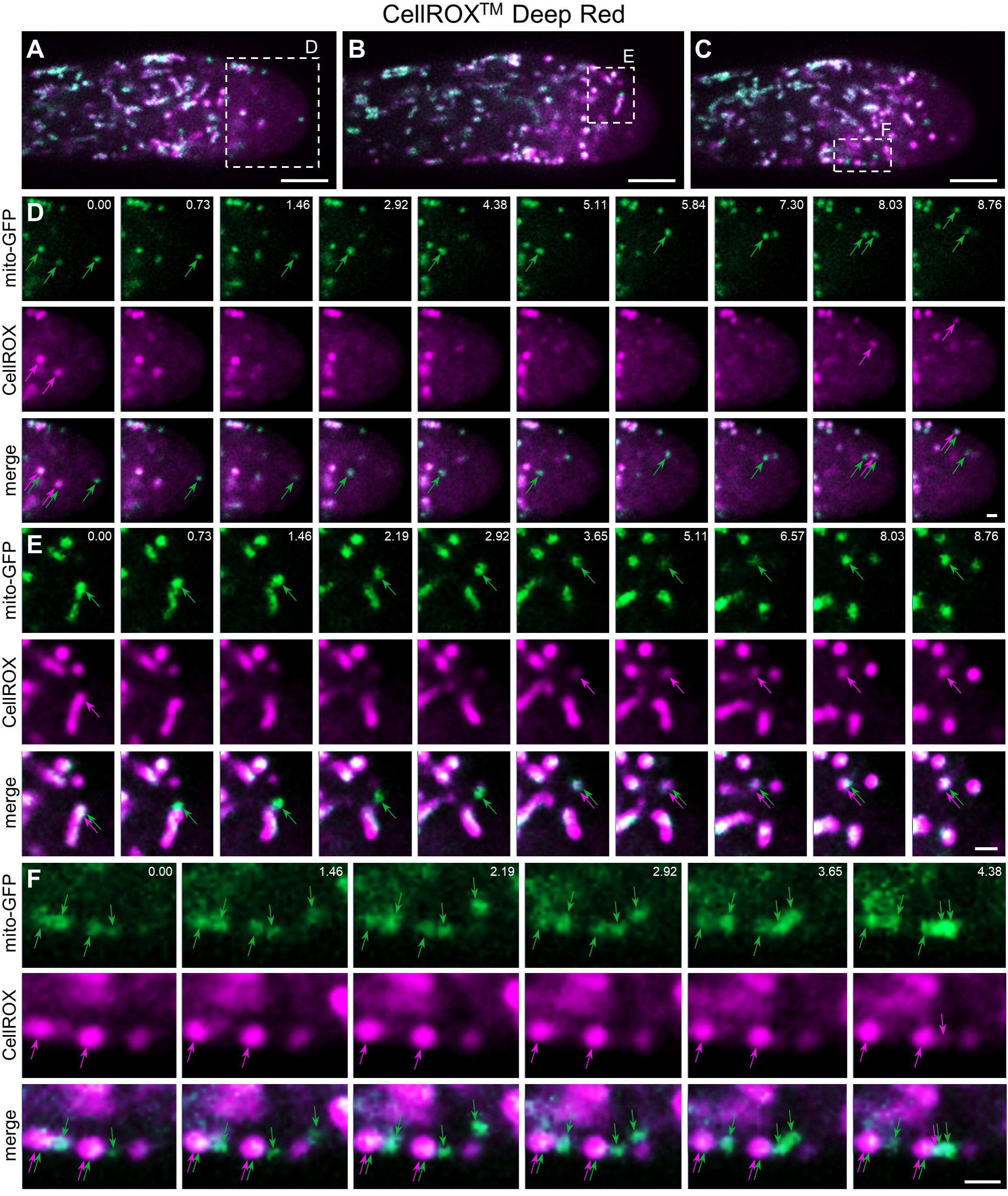
Single particle dynamic colocalization analysis of GFP-labelled mitochondria with ROS-accumulating compartments after staining with CellROX™ Deep Red in root hairs. **A-C.** Three different time points from time­ lapsed video-recordings of growing root hairs containing GFP-tagged mitochondria (in green) and ROS-accumulating compartments (in magenta) after CellROX™ Deep Red staining. **D.** Video-sequence captured from dashed box in **A** showing mitochondria (green arrows) and ROS-accumulating compartments (magenta arrows) colocalized (double arrows in merge), while small individual mitochondria in the clear zone are still free of ROS (green arrows only in merge). **E.** Video-sequence captured from dashed box in **B** showing small mitochondrion negative for ROS staining (green arrow) independent from larger mitochondria containing ROS (double arrows in merge). **F.** Video-sequence captured from dashed box in **C** showing large round and immobile mitochondria containing ROS (double arrows in merge) and small mobile individual mitochondria free of ROS (green arrows only in merge). Time sequences (in seconds) are 940 shown in mito-GFP channels **(D-F)**. Scale bar = 5 μm **(A-C)**, 1 μm **(D-F)**.

### Pattern of ROS labeling with Amplex^TM^ Red in root hairs

Staining pattern and fluorescence intensity distribution within the apex of growing root hairs in Col-0 after staining with Amplex^TM^ Red revealed a prominent signal at the root hair periphery, indicating the cell wall (apoplastic) localization (Figure 4A-C,J). We decided to proof this localization pattern using transgenic line carrying GFP-RHD2, a GFP-tagged AtRBOHC/RHD2, which is localized in early endosomes/TGN and apical plasma membrane in root hairs (Kuběnová et al., 2022). Staining of GFP-RHD2 line for ROS with Amplex^TM^ Red and treatment with 300 µmol·L^−1^ mannitol showed protoplast detachment from the cell wall in root hairs by plasmolysis. The surface of detached protoplast was visualized by GFP-RHD2 located in the plasma membrane. Amplex^TM^ Red-positive surface signal appeared at the remaining apoplast upon protoplast detachment caused by plasmolysis (Figure 7A-C), thus supporting the cell wall localization. To support the cell wall-specific detection of ROS by Amplex^TM^ Red, growing root hairs were double-labeled with cell wall-specific dye Calcofluor White (staining preferentially cellulose) and Amplex^TM^ Red (Figure 7D-F). Qualitative colocalization (Figure 7G,I-K) and semi-quantitative profile measurement (Figure 7H,L-M) revealed overlapping colocalization patterns between Amplex^TM^ Red and Calcofluor White. This was further validated by Calcofluor White and Amplex^TM^ Red double labeling approach combined with root hair plasmolysis by 300 µmol·L^−1^ mannitol (Figure 7N-R).

**Figure 7.**
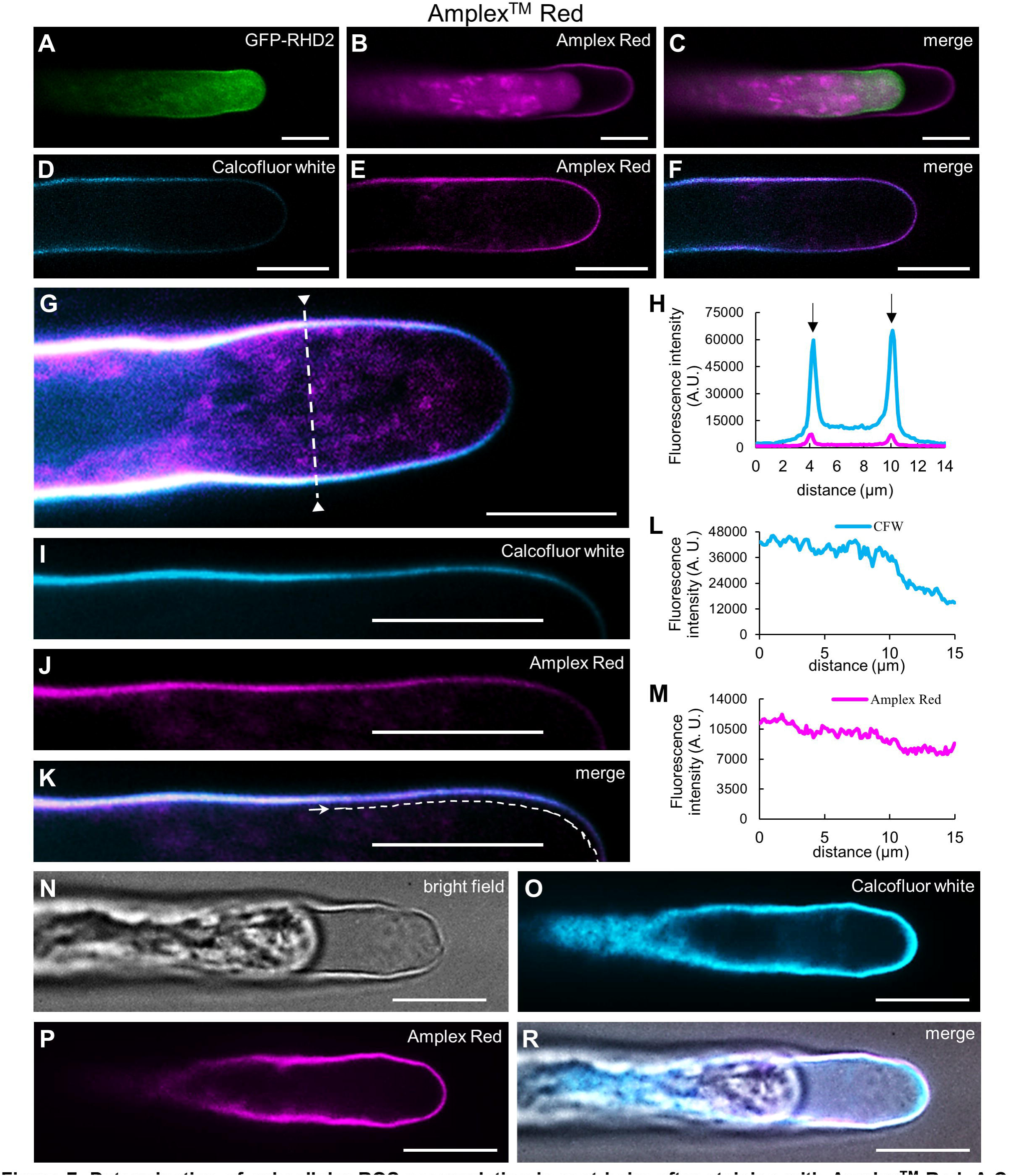
Determination of subcellular ROS accumulating in root hairs after staining with Amplex™ Red. **A-C.** Root hair of transgenic line carrying GFP-RHD2 plasmolyzed by 300 µmol·L-^1^ mannitol **(A)** that was before stained with Amplex™ Red **(B)**, shown in merged image **(C). D-F.** Growing root hair stained with Calcofluor White **(D)** that was after stained with Amplex™ Red **(E)**, shown in merged image **(F). G,H.** Colocalization of CalcofluorWhite and Amplex™ Red signal in sequentially stained root hair **(G)** and fluorescence intensity profile **(H)** of Calcofluor White (turquoise line) and Amplex^TM^ Red (magenta line) fluorescence signals along the interrupted white lines across the root hair shown in **(G). 1-M.** Colocalization of Calcofluor White **(I)** and Amplex^TM^ Red **(J)** signal in sequentially stained root hair **(K)** in the cell wall from the tip to subapical region and fluorescence intensity profile of Calcofluor White **(L)** and Amplex^TM^ Red **(M)** fluorescence signals along the interrupted white lines in **(K). N-R.** Root hair **(N)** stained with Calcofluor White **(0)** and plasmolyzed by 300 µmol·L-^1^ mannitol solution containing Amplex™ Red **(P)**, shown in merged image **(R).** Scale bar= 10 μm.

In addition to apoplastic signal, we also observed weak labeling of intracellular compartments by Amplex^TM^ Red in growing root hairs (Figure 7B,G). This was supported also by a pseudo-color-coded visualization of signal distribution (Figure 4A-C). Subsequently, we have used the colocalization analysis to characterize the nature of intracellular ROS-positive compartments in root hairs after Amplex^TM^ Red staining. GFP-RHD2 in moving vesicular compartments (Figure 8A) did not colocalize with ROS-positive compartments (Figure 8B,C). Early endosomal/TGN compartments marked by GFP-RabA1d (Figure 8D) did not show colocalization with ROS-positive compartments as well (Figure 8E,F). No colocalization between late endosomes marked by RabF2a-YFP (Figure 8G) and ROS-positive compartments (Figure 8H) was detected on merged images (Figure 8I). However, a GFP marker identifying mitochondria (Figure 8J) showed overlapping distribution pattern with ROS-positive compartments after Amplex^TM^ Red staining (Figure 8K), which was proved by fluorescence intensity profile measurements (Figure 8L). Colocalization analysis thus revealed that distinct motile compartments weakly stained by Amplex^TM^ Red in developing bulges and growing root hairs, are indeed mitochondria.

**Figure 8.**
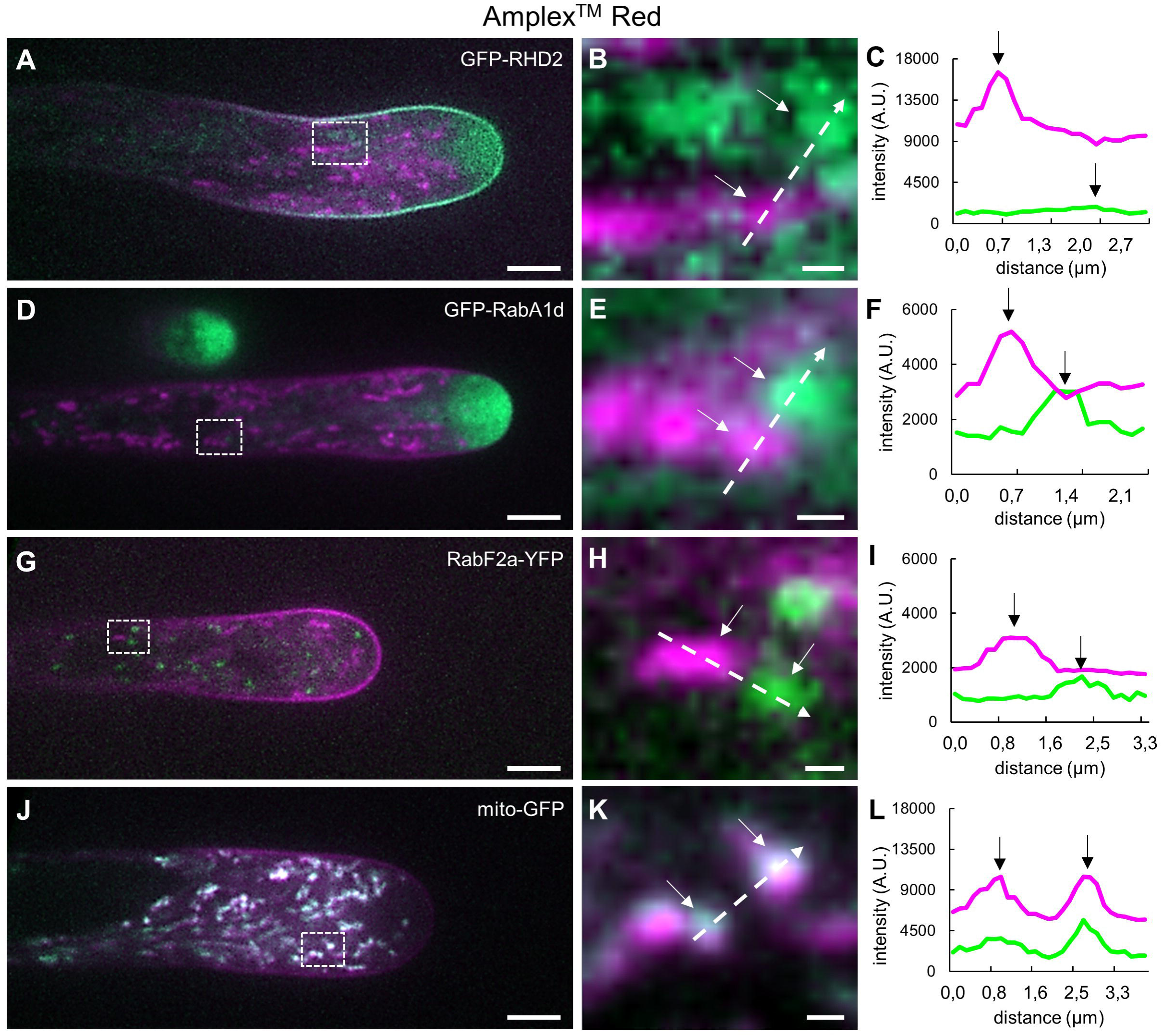
Colocalization analysis of subcellular molecular markers in root hairs with compartments accumulating ROS after staining with Amplex™ Red. **A-C.** Colocalization with transgenic line for visualization of vesicular compartments containing GFP-RHD2, a GFP-tagged AtRBOHC/RHD2. **D-F.** Colocalization with transgenic line carrying early endosomal marker GFP-RabA1d. **G-1.** Colocalization with transgenic line carrying late endosomal marker RabF2a-YFP. **J-L.** Colocalization with transgenic line carrying GFP-tagged mitochondria. Overview of root hairs carrying molecular markers (in green) and ROS­ accumulating compartments (in magenta) after Amplex™ Red staining **(A,D,G,J).** Detailed images of compartments containing carrying molecular markers and ROS **(B,E,H,K)** from the white boxes shown in **(A,D,G,J).** Fluorescence intensity profiles **(C,F,l,L)** of fluorescence signals of molecular markers (green lines) and ROS-containing compartments (magenta lines) along the interrupted white lines shown in **(B,E,H,K).** Arrows in **(B,C,E,F,H,l,K,L)** indicate the position of the compartments. Scale bar = 5 µm **(A,D,G,J)**, 0.5 µm **(B,E,H,K).**

### Spatial changes of ROS production and compartmentalization induced by external modulators

To further validate the specificity of probes in growing root hairs, we have used external modulators of ROS production and compartmentalization. Therefore, we tested the effect of ethylene, which is known to cause a ROS burst in root hairs. Considering physiological relevance of the experiment and efficient conversion of ACC to ethylene necessary for ROS production in trichoblasts, only low doses of ACC precursor has been used (according to Martin et al., 2022). Arabidopsis seedlings growing on media containing 0.7 µmol·L^-1^ ACC for 24h displayed a typical phenotype with shorter roots and a high density of root hairs (Supplemental Figure S8). Typical amount of ROS accumulation was observed in apical and subapical cytoplasm of growing root hairs after staining of Col-0 plants with CM-H_2_DCFDA (Figure 9A-C), which was significantly increased in root hairs pre-treated with ACC. Evaluation of images acquired at three different time points (0 min, 5 min and 10 min from the beginning of imaging) revealed enhanced fluorescence, which was distributed in whole apical and subapical parts of growing root hairs (Figure 9D-F). Measurement in root hair tips (Supplemental Figure S1B) confirmed considerable fluorescence intensity increases in ACC-treated root hairs (Figure 9G). Averaged root hair tip growth rate compared between control and ACC-treated Col-0 plants within the period of 10 min showed reduction caused by ACC treatment (Figure 9H). This is supported by kymographs showing velocity of the tip growth in representative individual root hairs and differences between control non-stained root hair (Figure 9I), CM-H_2_DCFDA-stained root hair (Figure 9J), and ACC-pretreated and CM-H_2_DCFDA-stained root hair (Figure 9K).

**Figure 9.**
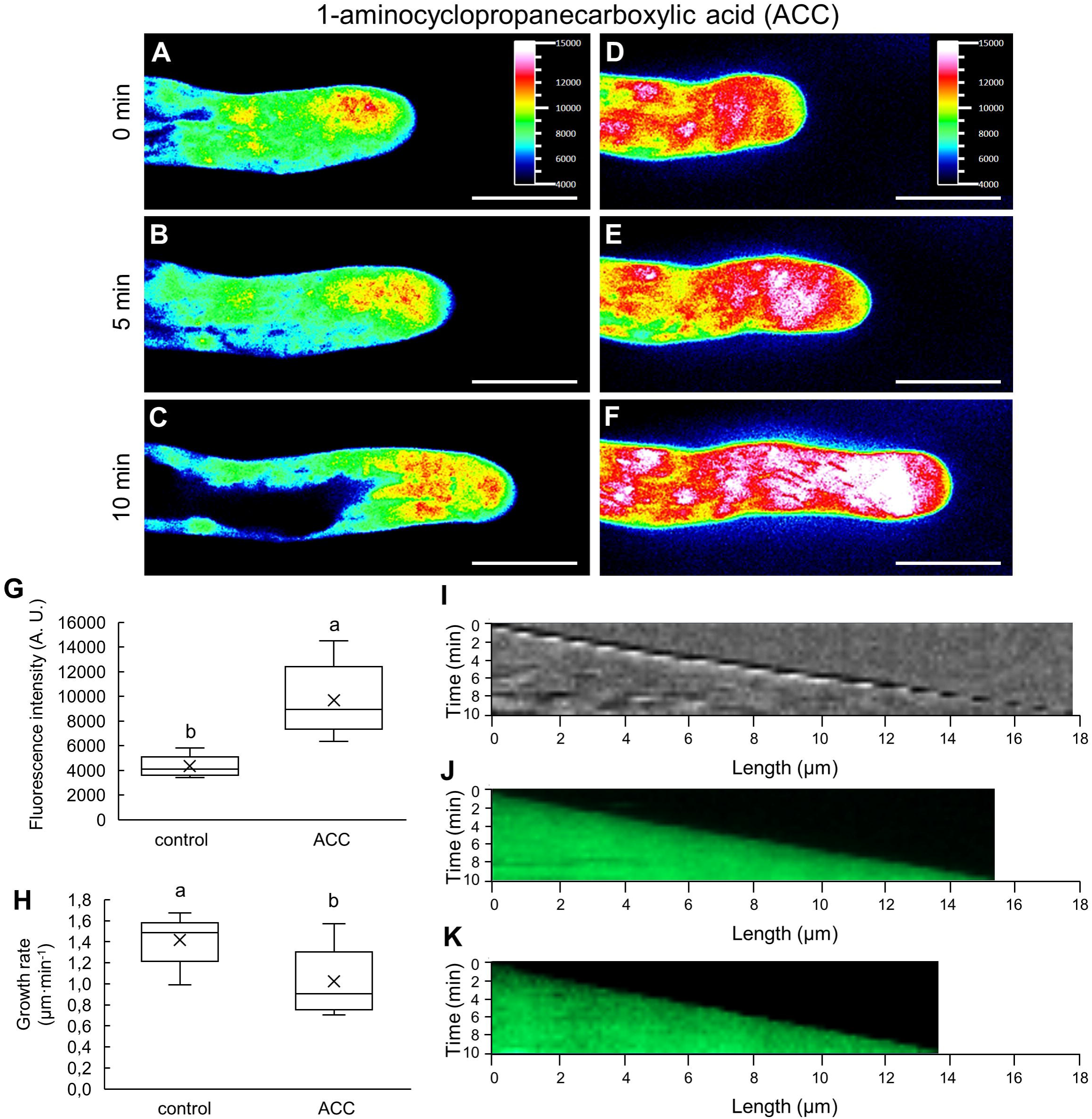
Accumulation of ROS in root hairs of Col-0 stained with CM-H_2_DCFDA after pre-treatment with the ethylene precursor ACC. **A-F.** Growing control root hair of Col-0 **(A-C)**, and root hair pre-treated with ACC **(D-F)** stained with CM-H_2_DCFDA for ROS localization and quantification. Root hairs were imaged at time points of O min **(A,D)**, 5 min **(B,E)** and 10 min **(C,F)** of growth. Fluorescence intensity distribution is visualized in a pseudo-color-coded scale, where black represents minimal intensity and white represents maximum intensity (insets in **A,D). G.** Semi-quantitative measurement of mean CM-H_2_DCFDA fluorescence intensity accumulation in tips of root hairs (measured area schematically illustrated in Supplemental Figure S1B) in control and ACC-treated Col-0 root hairs. **H.** Averaged root hair tip growth rate of control and ACC­ treated root hairs of Col-0 plants measured during imaging within the time period of 10 min. N = 9-10. Box plots display the first and third quartiles, split by the median; the crosses indicate the mean values; whiskers extend to include the max/min values. Lowercase letters indicate statistical significance between lines according to one-way ANOVA with Fisher’s LSD tests (P < 0.05). **1-K.** Kymographs presenting a velocity of root hair tip growth of control non-stained root hair (I), CM-H_2_DCFDA-stained root hair **(J)**, and ACC-treated and CM-H_2_DCFDA-stained root hair **(K).** Scale bar= 10 µm **(A-F).**

Mitochondria are very sensitive to any changes in their membranes potential, representing a driving force for the uptake of different cations. Valinomycin is a potent ionophore causing inner mitochondrial membrane depolarization and malfunctioning (Kolisek et al., 2003). We pre-treated Col-0 plants with valinomycin, and then analyzed CellROX^TM^ Deep Red-based staining and distribution of ROS in growing root hairs. Control root hairs of Col-0 stained with CellROX^TM^ Deep Red showed ROS-positive mitochondria distributed in cytoplasm mainly in the root hair subapical zone (Figure 10A-C; Supplemental Figure S9A-C). However, valinomycin pre-treatment caused removal of mitochondria-resident ROS fluorescence signal (Figure 10D-F; Supplemental Figure S9D-F). When we measured a relative distribution of pixels in subapical part of root hairs and displayed them in a graph according to their fluorescence intensity, we found a shallow peak of pixels with low fluorescence intensity and many counts of pixels with high fluorescence intensity in root hairs stained with CellROX^TM^ Deep Red alone (Figure 10G). Pixels with high fluorescence intensity in the graph (Figure 10G) from root hairs stained with CellROX^TM^ Deep Red alone are represented by white color in pseudo-color-coded images (Figure 10A-C). A similar analysis from valinomycin pre-treated root hairs revealed strong shifting of the curve to pixels with low fluorescence intensity, while high fluorescence intensity pixels were missing (Figure 10G). This is indicated by absence of white color in pseudo-color-coded images of valinomycin pre-treated root hairs (Figure 10D-F). Fluorescence pixel intensity was measured in area located in the subapical region of root hairs (Supplemental Figure S1D). Accordingly, measurement of mean fluorescence intensity accumulation in subapical part of root hairs confirmed a considerable reduction in valinomycin-pretreated root hairs (Figure 10H). Measurement during the recording time of 10 min revealed that averaged root hair tip growth rate of valinomycin-treated and CellROX^TM^ Deep Red-stained root hairs was significantly reduced in comparison to root hairs labeled by CellROX^TM^ Deep Red without valinomycin (Figure 10I). This was supported by kymographs showing velocity of the tip growth in representative individual root hairs and differences between control non-stained root hair (Figure 10J), CellROX^TM^ Deep Red-stained root hair (Figure 10K), and valinomycin-treated and CellROX^TM^ Deep Red-stained root hair (Figure 10L). The representative examples of root hair tip growth rate analysis by kymographs are provided for the control non-stained root hair (Supplemental Figure S10A), and root hairs stained by CM-H_2_DCFDA (Supplemental Figure S10B), CellROX^TM^ Deep Red (Supplemental Figure S10C), Amplex^TM^ Red (Supplemental Figure S10D) or FDA (Supplemental Figure S10E), along with averaged root hair tip growth rates (Supplemental Figure S10F).

**Figure 10.**
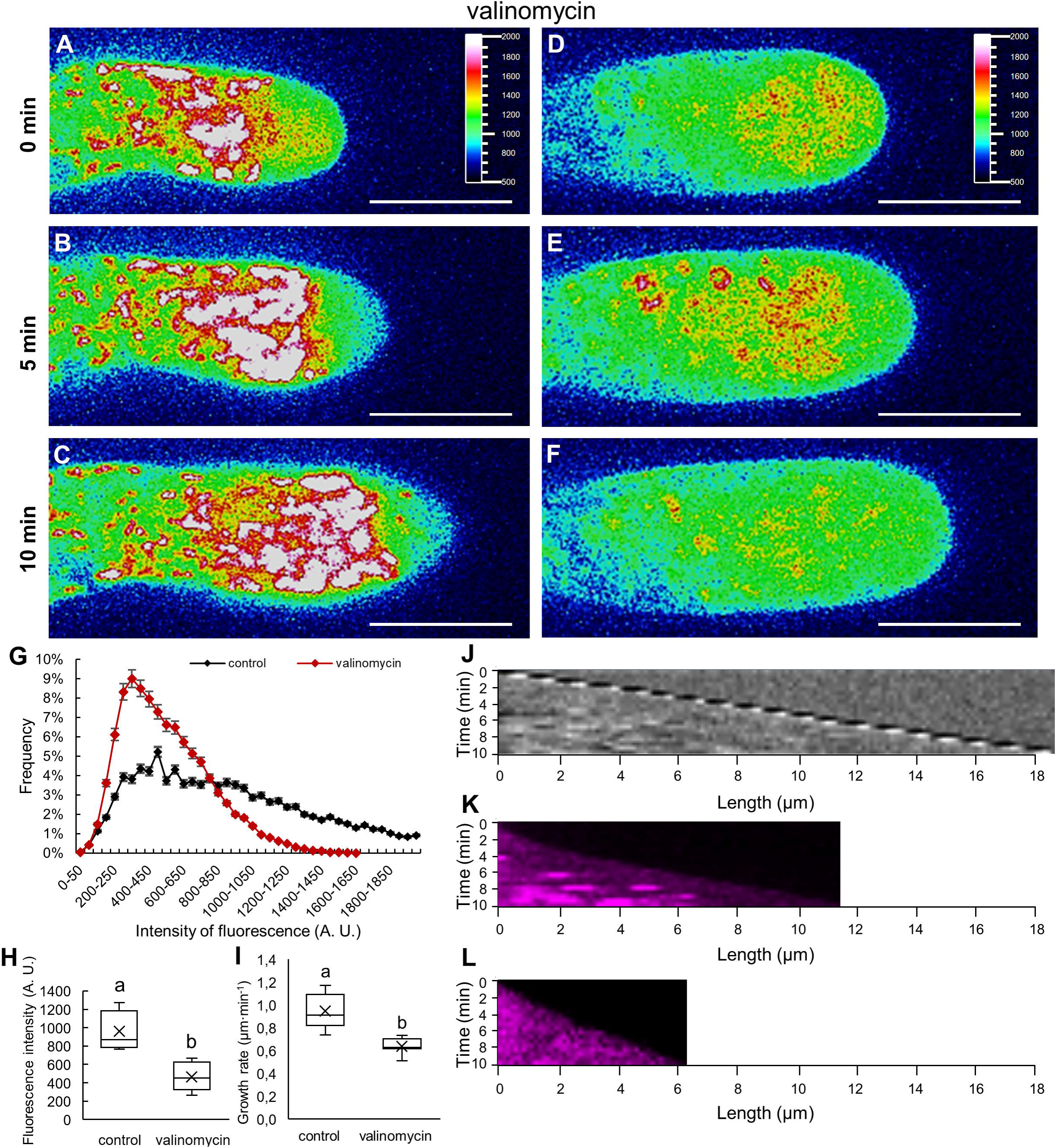
Redistribution of ROS stained with CellROX™ Deep Red in root hairs of Col-0 after pre-treatment with the mitochondrial ionophore valinomycin. **A-F.** Growing control root hair of Col-0 stained with CellROX™ Deep Red **(A-C)**, and root hair stained with CellROX™ Deep Red that was pre-treated with valinomycin **(D-F).** Root hairs were imaged at time points of O min **(A,D)**, 5 min **(B,E)** and 10 min **(C,F)** of growth. Fluorescence intensity distribution is visualized in a pseudo-color-coded scale, where black represents minimal intensity and white represents maximum intensity (insets in **A,D). G.** Relative distribution of pixels in apical part of root hairs according to their fluorescence intensity. Normalized root hair number was evaluated using 50 A.U. intervals distribution for control and valinomycin-treated root hairs. **H.** Semi-quantitative measurement of mean CellROX™ Deep Red fluorescence intensity accumulation in subapical part of root hairs (measured area schematically illustrated in Supplemental Figure S1D) in control and valinomycin-treated Col-0 root hairs. I. Averaged root hair tip growth rate of control and valinomycin­ treated root hairs of Col-0 plants measured during imaging within the time period of 10 min. N = 10. Box plots display the first and third quartiles, split by the median; the crosses indicate the mean values; whiskers extend to include the max/min values. Lowercase letters indicate statistical significance between lines according to one-way ANOVA with Fisher’s LSD tests (P < 0.05). **J­L.** Kymographs presenting a velocity of root hair tip growth of control non-stained root hair **(J)**, CellROX™ Deep Red-stained root hair **(K)**, and valinomycin-treated and CellROX™ Deep Red-stained root hair **(K).** Scale bar= 10 µm **(A-F).**

Altogether, experiments with external ROS modulators ACC and valinomycin confirmed the sensitivity of used fluorescent probes to specific changes in ROS production (induced by ACC) or subcellular compartmentalization (induced by valinomycin). The later experiments also indicate that physiological status of mitochondria might be important for ROS homeostasis during root hair tip growth.

## Discussion

Production, accumulation and localization of ROS in plant cells can be documented by different methods. Popular are small-molecule fluorescent probes that can assess ROS produced within cells, or even those released from cells. Because ROS represent a range of chemical oxygen species with different properties, investigating ROS in biological systems should selectively concern a particular ROS of interest. It is obvious that most of the ROS probes do not capture quantitatively intracellular ROS formed under investigation. In addition, the specific reaction of the probe with the ROS generated in cells may interfere with the cellular redox state and indeed, affect experimental results (Murphy et al., 2022). Small-molecule fluorescent probes are thus frequently used for ROS documentation. However, some limitations, such as selectivity, ability to quantify, linearity in response or some possible artifacts should be considered (Kalyanaraman et al., 2012). Fluorescent probe 2’,7’-dichlorodihydrofluorescein (DCFH), its diacetate form (CM-H_2_DCFDA), rapidly enters living cells and is used as an indicator of ROS in living cells. After its passive diffusion into the cytoplasm, its acetate groups are cleaved by esterases and chloromethyl groups react with glutathione and other thiols. It is subsequently oxidized to a highly fluorescent product 2’,7’-dichlorofluorescein (DCF), which is entrapped inside the cell. This allows long-term observation (Eruslanov and Kusmartsev, 2010). It is often used as a probe for the indirect detection of H_2_O_2_ and oxidative stress (Kalyanaraman et al., 2012; Winterbourn, 2014). However, oxidation to the fluorescent 2’,7’-dichlorofluorescein could be catalyzed by several ROS. Thus, this probe is not always considered specific for any particular ROS (Murphy et al., 2022). It is very important to assess physiologically relevant conditions for cells before ROS are detected, visualized and measured. In our system, undisturbed tip growth of root hairs, reflected by speed of growth and root hair morphology, secured biologically relevant conditions in our experiments (Supplemental Movie S1). In addition, we stained growing root hairs with a cell viability marker FDA, which served as a positive control for monitoring root hair viability and tip growth. Thus, viability and fitness of treated plants in the microscope were sustainably controlled. A slight increase in ROS levels during imaging of growing root hairs could be induced by the energy of lasers used in long-term observation (Khan et al., 2015). Light is absorbed by organic molecules such as flavins and they degrade in reaction with oxygen (Gorgidze et al., 1998; Hockberger et al., 1999; Rösner et al., 2016). This process then produces ROS, mainly O_2_ ^•−^ and hydroxyl radicals (Icha et al., 2017), which cause the formation of DCF (Eruslanov and Kusmartsev, 2010) and the subsequent increase in fluorescence intensity. Moreover, an increasing amount of ROS is typical during root hair elongation. Generation of ROS is important for the import of Ca^2+^ ions, which are required for apical cell expansion (Foreman et al., 2003; Carol et al., 2005). Through the control of root hair morphology, undisturbed tip growth, as well as fluorescent dyes loading, any difference in signal intensity among different dies, genotypes and recording time-points can be qualitatively and semi-quantitatively analyzed, with minimizing the side effects such as prolonged, delayed or unequal dye internalization during imaging.

The level of H_2_O_2_ in the apoplast can be determined by oxidizing peroxidase substrates, such as frequently used Amplex^TM^ Red. In addition to H_2_O_2_ produced in the apoplast, this probe can be sensitive to H_2_O_2_ released from cell, since H_2_O_2_ can cross plasma membrane directly or via aquaporins (Bienert and Chaumont, 2014). Importantly, the measurement could be a real balance between H_2_O_2_ being produced or removed by intracellular or apoplastic enzymes, and it may reflect the rate of diffusion not only inside, but also outside of the cell. Therefore, Amplex^TM^ Red with horseradish peroxidase can be used for the detection of H_2_O_2_ release from cells if other reducing agents or peroxidase substrates are absent (Murphy et al., 2022).

The process of ROS production in cells, in relation to their subcellular source, reflects either a passive production, where ROS are generated as by-products of metabolic pathways during photosynthesis and respiration, or active production by oxidases for signaling, such as RBOHs, respiratory burst oxidase homologues. The cellular sources of ROS production for signaling can thus be defined as extrinsic (in apoplast and/or cell wall), cytoplasmic and nuclear, or organellar (in chloroplasts, mitochondria, or peroxisomes) (Yao et al., 2002; Datt et al., 2003; Mittler et al., 2022). The NADPH oxidases are plasma membrane proteins accepting electrons from NADPH. This happens at the cytosolic side of the plasma membrane with the formation of molecular oxygen at the outer side, leading to the production of apoplastic O_2_^•−^ (Mittler, 2017; Smirnoff and Arnaud, 2019). Fluorescent probe 2’,7’-dichlorodihydrofluorescein-diacetate (H_2_DCFDA) that is particularly sensitive to H_2_O_2_, was used for estimation of time-course production and intracellular localization of ROS in Arabidopsis root tips exposed to salt stress. The non-fluorescent H_2_DCFDA become entrapped inside the cells by the surrounding plasma membrane once hydrolyzed by the intracellular esterases to H_2_DCF. After reaction with H_2_O_2_ fluorescent dichlorofluorescein (DCF) is produced, which was utilized for ROS localization in endosomes (Leshem et al., 2007). Essential processes such as vesicle trafficking and ROS formation, were compromised in the presence of LY294002, a PI3K-specific inhibitor. ROS generation inside endosomes and endocytosis were indispensable for tip growth. Preincubation of seedlings with LY294002 considerably reduced ROS levels in the root hairs, as estimated using the dye 2’,7’-dichlorodihydrofluorescein-diacetate (Lee et al., 2008). On the other hand, the use of the CellROX^TM^ Deep Red revealed that the signal was excluded from the apex of growing pollen tube. Instead, it was distributed throughout the body of the pollen tube. This analysis suggested that the tip-localized ROS signal does not represent O_2_^•−^ or hydroxyl radicals, but H_2_O_2_ in growing pollen tubes. Such results indicate that ROS generated from mitochondria are not major contributors to specific spatial distribution of H_2_O_2_ in the apex of growing pollen tubes (Do et al., 2019). Interestingly, both *rhd2-1* and *der1-3* mutants that differed in ROS accumulation in root hair bulges after CM-H_2_DCFDA and Amplex^TM^ Red staining did not show any difference in the pattern of ROS distribution after CellROX^TM^ Deep Red staining (Figure 3).

Root hair tip growth is regulated to secure the correct shape and size of bulges and elongating root hairs. Factors such as Ca^2+^, pH and ROS play an important role in this process (Mangano et al., 2016). In growing root hairs, ROS are produced by RHD2 (ROOT HAIR DEFECTIVE 2), a plasma membrane-resident NADPH oxidase. ROS regulate polar root hair expansion by activating calcium channels allowing Ca^2+^ uptake. Arabidopsis *rhd2-1* mutant is defective in Ca^2+^ uptake, and have impaired cell expansion, causing a phenotype of very short root hairs (Foreman et al., 2003). Our results show that the *rhd2-1* mutant line possesses a considerably lower amount of ROS in the bulge as analyzed by a CM-H_2_DCFDA fluorescence. These results are not surprising, as a mutation of the *RHD2* gene targets the NADPH oxidase AtRBOHC/RHD2, responsible for the production of ROS in root hairs (Foreman et al., 2003). It proves previous results that intracellular ROS levels in short root hairs of the *rhd2* mutant, detected using CM-H_2_DCFDA, were below the detection limit at pH 5 (Lee et al., 2008). Activity of AtRBOHC/RHD2 leads to the generation of apoplastic O_2_^•−^, from which H_2_O_2_ is subsequently produced by a reaction catalyzed by SOD (Mangano et al., 2016). Our data show a dramatic reduction in the formation of apoplastic ROS, particularly H_2_O_2_, that has been clearly documented by very low level of Amplex^TM^ Red staining. In turn, a reduction in the formation of apoplastic ROS can also result in reduced levels of H_2_O_2_ in the cytoplasm, which was in our results reflected by a lowering in fluorescence intensity after CM-H_2_DCFDA staining. The actin cytoskeleton participates in the polar elongation of cells, and is indispensable also for determining the tip growth in root hairs (Zheng et al., 2009; Wang et al., 2010; Pei et al., 2012). Together with actin, indispensable is the activity of actin-binding proteins. Actin-depolymerization factor (ADF) and profilin are accumulated in the bulge and in the growing tips of root hairs, where they are responsible for the dynamic remodeling of the filamentous actin network (Staiger et al., 1997; Braun et al., 1999). A high degree of actin dynamics in the bulge selectively concentrates endomembranes and leads to the direction of vesicular transport preferentially to this subcellular domain (Baluška et al., 2000). Mutagenesis of the *ACTIN2* gene led to the isolation of *der1-3* mutant line, which shows a phenotype of very short root hairs not able to elongate after a bulge formation (Ringli et al., 2002; Vaškebová et al., 2018). Since *der1-3* mutant is more resistant to oxidative stress (Kuběnová et al., 2021), it justifies our interest to determine ROS accumulation and distribution in bulges of this mutant. Determination of ROS related to bulge stage during the root hair formation process revealed, however, that ROS levels selectively detected by CM-H_2_DCFDA and Amplex^TM^ Red staining were not altered in *der1-3* mutant. Apart from striking differences in cytoplasmic and apoplastic ROS levels between wild-type and *rhd2-1*, clear differences were found also between *rhd2-1* and *der1-3*, suggesting principally different spatiotemporal distribution, production, delivery and utilization of ROS in these mutants.

The results show that the employed dyes have high sensitivity for ROS localization in different subcellular compartments and therefore, are suitable for high-resolution imaging of ROS specific distribution in living cells. Morphological and molecular identification of ROS-positive compartments in root hairs after the use of different dyes is thus important aspect of the study. Our colocalization analysis identified motile compartments labeled with CellROX^TM^ Deep Red as mitochondria. In addition, it provided novel biological insight into possible participation of mitochondria in the maintenance of subapical region of root hair. In particular, the clear zone of growing root hairs is usually free of larger organelles including mitochondria that only sparsely invade this region (Zheng et al., 2009), thus their appearance and behavior in the clear zone is rather enigmatic. Here we document presence of mitochondria in the clear zone of growing root hairs. Although these were rather unique events, mitochondria appeared there temporarily and were temporarily not stainable for ROS with CellROX^TM^ Deep Red. Importantly, such mitochondria were small, round and highly motile. They were negative to CellROX^TM^ Deep Red staining at the beginning, however, this labeling was replenished in them after short time. In contrast, larger and less mobile mitochondria were intensely stained for ROS with CellROX^TM^ Deep Red. Appearance of small ROS-free mitochondria was not observed in root hair basal parts. Hypothetically, they may represent pre-mature mitochondria created by fission from mother ones (Rose, 2021; Westrate et al., 2014), while ROS producing capacity is activated in them only after some time.

Based on high sensitivity of employed dyes, external modulation of ROS production or compartmentalization should be microscopically detectable and directly analyzed in growing root hairs. Plant gas hormone ethylene affects root growth and causes elevated production of ROS in trichoblasts (Martin et al., 2022). An ethylene precursor, ACC, used in a high concentration may directly alter several developmental processes without its own conversion to ethylene (Li et al., 2022). Therefore, we applied recommended low doses of ACC (Martin et al., 2022) to keep physiological relevance in our experiments. Qualitative and quantitative analysis of mean fluorescence intensity accumulation in tips of growing root hairs after staining with CM-H_2_DCFDA revealed a significant increase in ACC-treated root hairs. This experiment confirmed the sensitivity of the dye CM-H_2_DCFDA to the external modulation of ROS production in cytoplasm.

Rapid influx of different cations (e.g. Mg^2+^) to mitochondria is driven by the mitochondrial membranes potential. Upon addition of ionophore valinomycin, the depolarization of mitochondrial membranes leads to drastic reduction of Mg^2+^ influx to mitochondria (Kolisek et al., 2003). We used this approach to depolarize mitochondrial membranes potential by valinomycin in root hairs. Roots of Col-0 plants were pre-treated with valinomycin and stained with CellROX^TM^ Deep Red. Compared to non-treated Col-0 plants, where root hairs contained mitochondria stained with CellROX^TM^ Deep Red, such specific staining pattern disappeared after pre-treatment with valinomycin. This result indicates that depolarization of mitochondrial membranes potential by valinomycin likely prevents CellROX^TM^ Deep Red oxidation inside mitochondria, which might be impaired in ROS production. This may suggest that in control root hairs, CellROX^TM^ Deep Red internalization to fully-functional mitochondria is needed for its ROS-dependent oxidation.

We have used a semi-quantitative measurement approach, which is compatible with high-resolution live-cell imaging microscopy. A quantitative approach would also be feasible, considering the biochemical principle of the reactions required for fluorescence detection. However, the reaction of the probe with the ROS generated in cells may interfere with the cellular redox state. As a consequence, the ROS equilibrium can be changed by reducing the number of molecules to be detected due to their use for a probe oxidation (Kalyanaraman et al., 2012). For example, the Amplex^TM^ Red is an oxidizing substrate for peroxidases in the apoplast, and can be used with horseradish peroxidase for detection of H_2_O_2_ release from cells, but only if other reducing agents or peroxidase substrates are absent (Murphy et al., 2022). Therefore, the influence of employed dyes on ROS homeostasis in root hairs may affect directly or indirectly root hair tip growth rate. Indeed, our data show that averaged root hair tip growth rate was highest in the untreated root hairs, while it was decreased after FDA, ACC, CM-H_2_DCFDA, and Amplex^TM^ Red application. The growth reduction was considerable in CellROX^TM^ Deep Red-stained root hairs, and the highest one was recorded in valinomycin-treated and CellROX^TM^ Deep Red-stained root hairs (Figure 9,10; Supplemental Figure S10). Therefore, the possibility of partially affecting root hair physiology by the employed dyes through their scavenging potential to ROS should be taken into account.

In conclusion, our high-resolution live-cell imaging provides a dynamic mapping of ROS in cytoplasm, mitochondria and apoplast, highlighting subcellular location of the ROS production and dynamics in tip-growing root hairs (Figure 11). The data show a sensitivity of CM-H_2_DCFDA to ROS in cytoplasm, CellROX^TM^ Deep Red to ROS in mitochondria, and root hair-specific sensitivity of Amplex^TM^ Red to ROS in apoplast and mitochondria. The high sensitivity of CellROX^TM^ Deep Red combined with high spatiotemporal resolution imaging method revealed small, highly motile pre-mature mitochondria in the growing tip of root hairs that are free of ROS at the early stages of their formation. We also provide characteristic spatial changes in ROS production and compartmentalization that were induced by external ROS modulators, either by inducer, ethylene precursor ACC, or by membrane potential depolarization agent, ionophore valinomycin. Thus, this study also shows physiological characterization of ROS probes employed herein. Moreover, subcellular localization and quantification of ROS production in *rhd2-1* and *der1-3* root hair mutants, differences between them, and their distinct performance from corresponding wild-types shed new light on the relationship between different ROS sources in the root hair tip growth mechanism requiring both RBOHC and actin cytoskeleton. Overall, this study establishes fluorescent ROS probes, CM-H_2_DCFDA, CellROX^TM^ Deep Red and Amplex^TM^ Red, for functional characterization of spatiotemporal mode of ROS production, delivery and utilization in Arabidopsis wild-type bulges and root hairs, but also in diverse root hair mutants.

**Figure 11.**
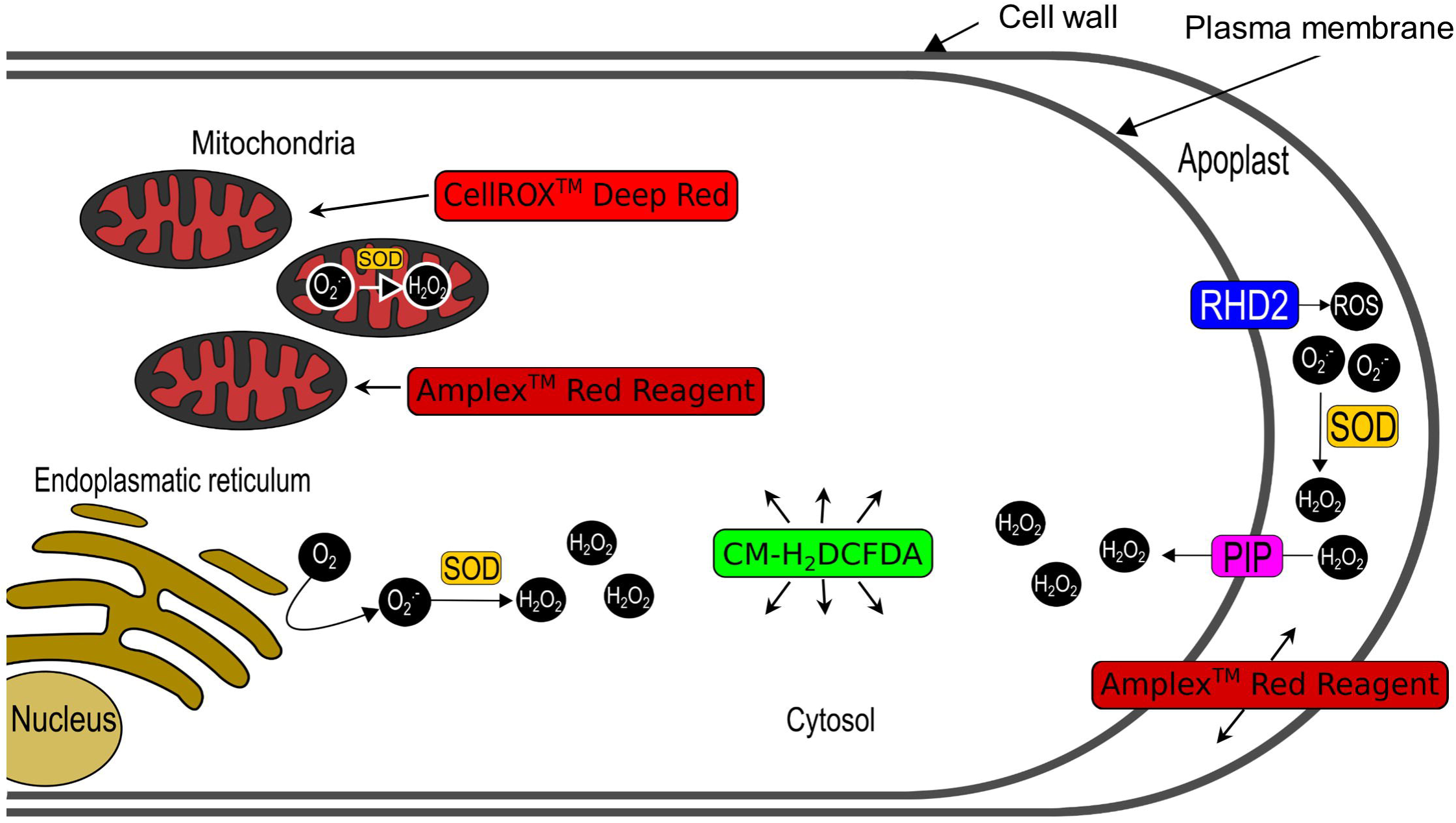
Model of subcellular ROS localization in root hairs with CM-H_2_DCFDA, CellROX™ Deep Red and Amplex™ Red. CM-H_2_DCFDA represents a sensitive probe for ROS in the cytoplasm. CellROX™ Deep Red localizes specifically ROS in mitochondria. Amplex^TM^ Red serves as a highly sensitive marker for root hair-specific apoplastic ROS. In addition, root hairs are able to internalize Amplex™ Red, which sensitively detects ROS in mitochondria.

## Materials and methods

### Plant material and cultivation in vitro

Experiments were performed with plants of *Arabidopsis thaliana* wild types of the Col-0 and C24 ecotypes, *rhd2-1* mutant (Foreman et al., 2003) and *der1-3* mutant (Ringli et al., 2002) and stably transformed Arabidopsis lines carrying construct GFP-RHD2 for NADPH oxidase type C (Takeda et al., 2008), GFP-RabA1d for early endosomes/TGN (Ovečka et al., 2010; Berson et al., 2014; von Wangenheim et al., 2016), RabF2a-YFP for late endosomes (von Wangenheim et al., 2016), and the GFP-tagged mitochondria-targeting sequence of the N-terminus of the F1-ATPase g-subunit (Niwa et al., 1999). After surface-sterilization, the seeds of control, mutant and transgenic lines were planted on half-strength Murasige and Skoog (MS) medium (Murashige and Skoog, 1962) without vitamins, solidified with 0.6% (w/v) gellan gum (Alfa Aesar, Thermo Fisher Scientific, Heysham, UK), the pH was adjusted to 5.7. To synchronize seed germination, Petri dishes were stratified for 3 days at 4°C and then cultivated vertically in an environmental chamber at 21°C, 70% humidity, and 16-h/8-h light/dark cycle. The illumination intensity was 130 µmol·m^−2^s^−1^.

### Application of fluorescent probes for ROS detection

CM-H_2_DCFDA, CellROX™ Deep Red and Amplex™ Red reagents (Thermo Fisher Scientific, Waltham, MA, USA) were used as vital probes for ROS detection in growing root hairs. At least in five independent plants per treatment, CM-H_2_DCFDA (5 µmol·L^−1^), CellROX™ Deep Red (10 µmol·L^−1^), and Amplex™ Red (1 µmol·L^−1^) in modified liquid MS medium (Ovečka et al., 2014) were individually applied by perfusion directly to plantlets in the microscopic chamber. In total, 60 µL of the modified liquid MS medium with CM-H_2_DCFDA, CellROX™ Deep Red, or Amplex™ Red was applied in 6 separate steps of 10 µL each. Plants were then observed using a spinning disk microscope. The root hairs were scanned at different times after perfusion of the probes. Several hairs were usually scanned consecutively, with at least 10 min pauses between scans to avoid induction of fluorescence caused by overloaded laser exposure.

### Plasmolysis of root hairs

Transgenic plants carrying GFP-RHD2, 2-to 3-day-old were transferred to microscopic chambers containing liquid MS medium modified according to Ovečka et al. (2005, 2014). After manipulation with plants during sample preparation, the subsequent stabilization period for 24 h allowed undisturbed growth of the root and the formation of new root hairs. The plasmolysis was induced with 300 mmol·L^−1^ mannitol (Sigma-Aldrich, St Louis, MO, USA) by perfusion methods. Amplex™ Red Reagent (1 µmol·L^−1^) was added to the mannitol solution. Total volume of solution was 100 µl divided into 10 steps of 10 µl applications. Plasmolyzed root hairs were immediately observed by Airyscan microscope.

### Epifluorescence microscopy

Plantlets of Col-0 plants (prepared in the microscopic chambers) were observed using an epifluorescence microscope Axio Imager M2 equipped with DIC optics and epifluorescence metal halide source illuminator HXP 120 V (Zeiss, Oberkochen, Germany), and analyzed using Zeiss ZEN 2012 Blue software (Zeiss, Germany). Root tips were labeled with fluorescent probes for ROS detection (CM-H_2_DCFDA, CellROX™ Deep Red and Amplex™ Red). Imaging was performed with N-Achroplan 5×/0.15 NA dry objective and documented with a Zeiss AxioCam ICm1 camera. A filter set providing a wavelength of 450-490 nm for the excitation and 515-565 nm for the emission at 200 ms exposure time was used to visualize the CM-H_2_DCFDA signal. For the CellROX™ Deep Red and Amplex™ Red, the filter set provided an excitation wavelength of 533-558 nm and an emission wavelength of 570-640 nm. The exposure time was set up to 1 s for both probes. The image scaling for all signals was 0.93 × 0.93 µm in x × y dimensions, and 15.12 µm in the z dimension. Final images were processed by orthogonal projection with weighted average function.

### Spinning disk (SD) microscopy

Samples were prepared according to Ovečka et al. (2005, 2014), by transferring 2-to 3-d-old plants to microscopic chambers containing modified liquid MS medium. As the root hairs are very sensitive to stress during the manipulation, overnight cultivation for plant stabilization and undisturbed root growth and the formation of new root hairs was necessary. For quantitative and colocalization analyses, a Cell Observer SD Axio Observer Z1 spinning disk microscope (Carl Zeiss,Germany) was used, equipped with alpha Plan-Apochromat 100X/1.57 NA DIC Korr oil immersion objective (Carl Zeiss, Germany). The samples were imaged using an excitation laser line of 405 nm and emission filter BP450/50 for Calcofluor White signal detection, excitation laser line of 488 nm and emission filter BP525/50 for CM-H_2_DCFDA, FDA and GFP signal detection, excitation laser line of 514 nm, and emission filter BP535/30 for YFP signal detection, excitation laser line of 639 nm and emission filter BP690/50 for CellROX^TM^ Deep Red signal detection, and excitation laser line of 561 nm and emission filter BP629/62 for the Amplex^TM^ Red signal detection. The excitation laser power level for all lasers used was set up to 50%, and the image scaling for all channels was 0.133 X 0.133 µm in x X y dimensions, and with the z dimension 0.50 µm for the Calcofluor White signal, 0.63 µm for the GFP, FDA and CM-H_2_DCFDA signal, 0.66 µm for the YFP channel, 0.75 µm for the CellROX^TM^ Deep Red and Amplex^TM^ Red signal. Images were acquired sequentially or simultaneously with two Evolve 512 EMCCD cameras (Photometrics) with an exposure time of 700 ms. The samples were scanned with 700 ms of exposure time for GFP, YFP and CM-H_2_DCFDA signal, with 50 ms of exposure time for CellROX^TM^ Deep Red and Calcofluor White signal, and with 500 ms of exposure time for Amplex^TM^ Red signal (for quantitative analysis) or using a camera-streaming mode (for colocalization studies).

### Airyscan confocal laser scanning microscopy (Airyscan CLSM)

Root hairs for ROS staining after plasmolysis were observed using a confocal laser scanning microscope LSM880 equipped with Airyscan (Carl Zeiss, Germany). Image acquisition was performed with a 20×/0.8 NA dry Plan-Apochromat objective (Carl Zeiss, Germany). The samples were imaged with an excitation laser line of 488 nm and BP420-480 + BP495-550 emission filters for GFP detection and an excitation laser line of 561 nm and BP420-480 + BP600-650 emission filters for Amplex^TM^ Red detection. The laser power did not exceed 2% for GFP and 1% for Amplex^TM^ Red of the available laser intensity range. The samples were scanned with 700 ms of exposure time for both signals using a 32 GaAsP detector. Pixel dwell time was set up to 1.6 µs, and with default settings of the gain level the image scaling was set up to 0.07 × 0.07 µm (x × y).

### ROS induction

For ROS induction in root hairs, 1-aminocyclopropanecarboxylic acid (ACC), an ethylene precursor, was used according to Martin et al. (2022). Two-day-old seedlings of *A. thaliana* Col-0, previously cultured on the control medium, were carefully transferred to the medium containing 0.7 µmol·L^-1^ ACC. After transfer to medium with ACC, plantlets were cultured *in vitro* overnight, vertically in a phytotron at 21 °C, 70 % humidity and a photoperiod of 16/8 h light/dark (illumination intensity 130 µmol·m^−2^·s^−1^). The next day, samples were prepared as described above by transferring to microscopic chambers containing modified liquid MS medium (Ovečka et al., 2014), containing 0.7 µmol·L^-1^ ACC. The prepared chambers were cultured overnight in a phytotron at 21 °C, 70 % humidity and a photoperiod of 16/8h light/dark (illumination intensity 130 µmol·m^−2^s^−1^). Next day, the CM-H_2_DCFDA dye (5 µmol·L^-1^) was applied to prepared chambers by perfusion and examined at the microscope.

### Inhibition of mitochondria

Valinomycin (Sigma-Aldrich, St Louis, MO, USA), applied directly to microscopic chambers by perfusion, at a final concentration of 5 µmol·L^-1^ (Kolisek et al., 2003), was used to induce membrane potential depolarization in mitochondria. Subsequently, the prepared chambers with plants and applied valinomycin were incubated for 30 minutes and then the CellROX^TM^ Deep Red dye (10 µmol·L^-1^) was applied by perfusion and treated plants in chambers were examined at the microscope.

### Cell wall staining

Calcofluor White stain binding to cellulose in plant cell walls, dissolved in a modified culture medium, was applied to the plants in microscopic chambers using the perfusion method in a final concentration of 100 µmol·L^-1^. The chambers were then incubated in the dark for 90 min, according to Bidhendi et al. (2020). Then root hair plasmolysis was induced using 300 mmol·L^−1^ D-mannitol. Amplex™ Red (1 µmol·L^−1^) was added directly to the solution with mannitol. For experiments without plasmolysis, after incubation with Calcofluor White (100 µmol·L^-1^) only Amplex^TM^ Red dye was applied to plants in microscopic chambers.

### Data analysis and measurements

The signal intensity was analyzed using a spinning disk microscope. The root hairs were scanned for 10 min at 30 s intervals and sorted into groups according to developmental stages. For Col-0 and C24 lines at the stages of bulge, short root hair (10-200 µm) and long root hair (200+ µm), in *der1-3* and *rhd2-1* mutant lines at the bulge stage. Zen 3.3 blue edition software (Carl Zeiss, Germany) was used to obtain profiles for quantitative signal intensity distribution of different fluorescent probes in growing root hairs. Fluorescence intensity distribution was measured in single optical section, along the profile in a 10 µm segment from the center of the root hair to its tip (Supplemental Figure S1A), or as the mean fluorescence intensity in area of the tip (Supplemental Figure S1B). For the Amplex^TM^ Red probe, an area including only the cell wall was measured (Supplemental Figure S1C). Distribution of the CellROX^TM^ Deep Red before and after valinomycin treatment was measured in area located in the subapical region of the root hair (Supplemental Figure S1D). Measurements were always performed at the start (0 min), in the middle (5 min) and at the end (10 min) of imaging. The mean fluorescence intensity in area of the tip (Supplemental Figure S1B) was always analyzed in the middle of the imaging time (5 min). In vacuolated bulges, the area was adjusted individually to not include vacuoles to the measurement.

Acquired data were processed in Microsoft Excel. Individually, the mean background intensities were subtracted from the intensities in the segment. Root hairs with abnormal intensities (exceeding double intensity over the average) were discarded from the statistics. All charts were done in Microsoft Excel software and statistical analyses were obtained using STATISTICA 13.4 (StatSoft) software using analysis of variance (ANOVA) and subsequent Fisher’s LSD tests (P < 0.05). The colocalization studies of fluorescence signals were analyzed using a spinning disk microscope by simultaneous signal acquisition with two independent Evolve 512 EMCCD cameras. The colocalization was analyzed in transgenic lines GFP-RHD2, mito-GFP, GFP-RabA1d, and RabF2a-YFP in combination with the fluorescent probes for ROS detection described above. Profiles for colocalization were generated using Zen Blue 2014 software (Carl Zeiss, Germany) and graphically edited in Microsoft Excel.

## Supplemental Data

**Supplemental Figure S1.** Topology of the quantitative signal intensity measurement in bulges and growing root hairs.

**Supplemental Figure S2.** Distribution of ROS in Col-0 wild-type, *rhd2-1* and *der1-3* mutants after staining with CM-H_2_DCFDA.

**Supplemental Figure S3.** Fluorescence intensity measurements in apical parts of bulges and root hairs after ROS staining with CM-H_2_DCFDA.

**Supplemental Figure S4.** Distribution of ROS in Col-0 wild-type, *rhd2-1* and *der1-3* mutants after staining with CellROX^TM^ Deep Red.

**Supplemental Figure S5.** Fluorescence intensity measurements in apical parts of bulges and root hairs after ROS staining with CellROX^TM^ Deep Red.

**Supplemental Figure S6.** Distribution of ROS in Col-0 wild-type, *rhd2-1* and *der1-3* mutants after staining with Amplex^TM^ Red.

**Supplemental Figure S7.** Fluorescence intensity measurements in apical parts of bulges and root hairs after ROS staining with Amplex^TM^ Red.

**Supplemental Figure S8.** Phenotype of Col-0 wild-type seedlings treated with the ethylene precursor ACC.

**Supplemental Figure S9.** Distribution of ROS stained with CellROX^TM^ Deep Red in root hairs of Col-0 after pre-treatment with the mitochondrial ionophore valinomycin.

**Supplemental Figure S10.** Speed of the root hair tip growth analyzed by kymographs.

**Supplemental Movie S1_Col-0_control_root_hair.** Growing unstained root hairs.

**Supplemental Movie S2_Col-0_CM-H2DCFDA_bulge.** Developing bulge of Col-0 wild-type imaged with CM-H_2_DCFDA.

**Supplemental Movie S3_Col-0_CM-H2DCFDA_root_hair.** Growing root hair of Col-0 wild-type imaged with CM-H_2_DCFDA.

**Supplemental Movie S4_rhd2_CM-H2DCFDA_bulge_measurement.** Developing bulge of *rhd2-1* imaged with CM-H_2_DCFDA and profile measurement of fluorescence intensity.

**Supplemental Movie S5_der1_CM-H2DCFDA_bulge_measurement.** Developing bulge of *der1-3* imaged with CM-H_2_DCFDA and profile measurement of fluorescence intensity.

**Supplemental Movie S6_Col-0_CM-H2DCFDA_bulge_measurement.** Developing bulge of Col-0 wild-type imaged with CM-H_2_DCFDA and profile measurement of fluorescence intensity.

**Supplemental Movie S7_Col-0_CM-H2DCFDA_root_hair_measurement.** Growing root hair of Col-0 wild-type imaged with CM-H_2_DCFDA and profile measurement of fluorescence intensity.

**Supplemental Movie S8_Col-0_CellROXTM Deep Red_bulge.** Developing bulge of Col-0 wild-type imaged with CellROX^TM^ Deep Red.

**Supplemental Movie S9_Col-0_CellROXTM Deep Red_root_hair.** Growing root hair of Col-0 wild-type imaged with CellROX^TM^ Deep Red.

**Supplemental Movie S10_rhd2_CellROXTM Deep Red_bulge_measurement.** Developing bulge of *rhd2-1* imaged with CellROX^TM^ Deep Red and profile measurement of fluorescence intensity.

**Supplemental Movie S11_der1_CellROXTM Deep Red_bulge_measurement.** Developing bulge of *der1-3* imaged with CellROX^TM^ Deep Red and profile measurement of fluorescence intensity.

**Supplemental Movie S12_Col-0_CellROXTM Deep Red_bulge_measurement.** Developing bulge of Col-0 wild-type imaged with CellROXTM Deep Red and profile measurement of fluorescence intensity.

**Supplemental Movie S13_Col-0_CellROXTM Deep Red_root_hair_measurement.** Growing root hair of Col-0 wild-type imaged with CellROX^TM^ Deep Red and profile measurement of fluorescence intensity.

**Supplemental Movie S14_Col-0_AmplexTM Red_bulge.** Developing bulge of Col-0 wild-type imaged with Amplex^TM^ Red.

**Supplemental Movie S15_Col-0_AmplexTM Red_root_hair.** Growing root hair of Col-0 wild-type imaged with Amplex^TM^ Red.

**Supplemental Movie S16_rhd2_AmplexTM Red_bulge_measurement.** Developing bulge of *rhd2-1* imaged with Amplex^TM^ Red and profile measurement of fluorescence intensity.

**Supplemental Movie S17_der1_AmplexTM Red_bulge_measurement.** Developing bulge of *der1-3* imaged with Amplex^TM^ Red and profile measurement of fluorescence intensity.

**Supplemental Movie S18_Col-0_AmplexTM Red_bulge_measurement.** Developing bulge of Col-0 wild-type imaged with Amplex^TM^ Red and profile measurement of fluorescence intensity.

**Supplemental Movie S19_Col-0_AmplexTM Red_root_hair_measurement.** Growing root hair of Col-0 wild-type imaged with Amplex^TM^ Red and profile measurement of fluorescence intensity.

## Authors’ contributions

LK and JH performed experiments, microscopic data acquisition and analysis, semi-quantitative evaluation and statistical analyses. PD analyzed data from fluorescent probe labeling. JŠ and MO contributed to the experimental plan and data interpretation. MO wrote the manuscript with input from all co-authors. JŠ provided infrastructure and secured funding.

## Data Availability

Data that support the findings of this study are available from the corresponding author upon reasonable request.

