## Supplemental Figures S1-S10 for "Spatiotemporal distribution of ROS production, delivery and utilization in Arabidopsis root hairs"

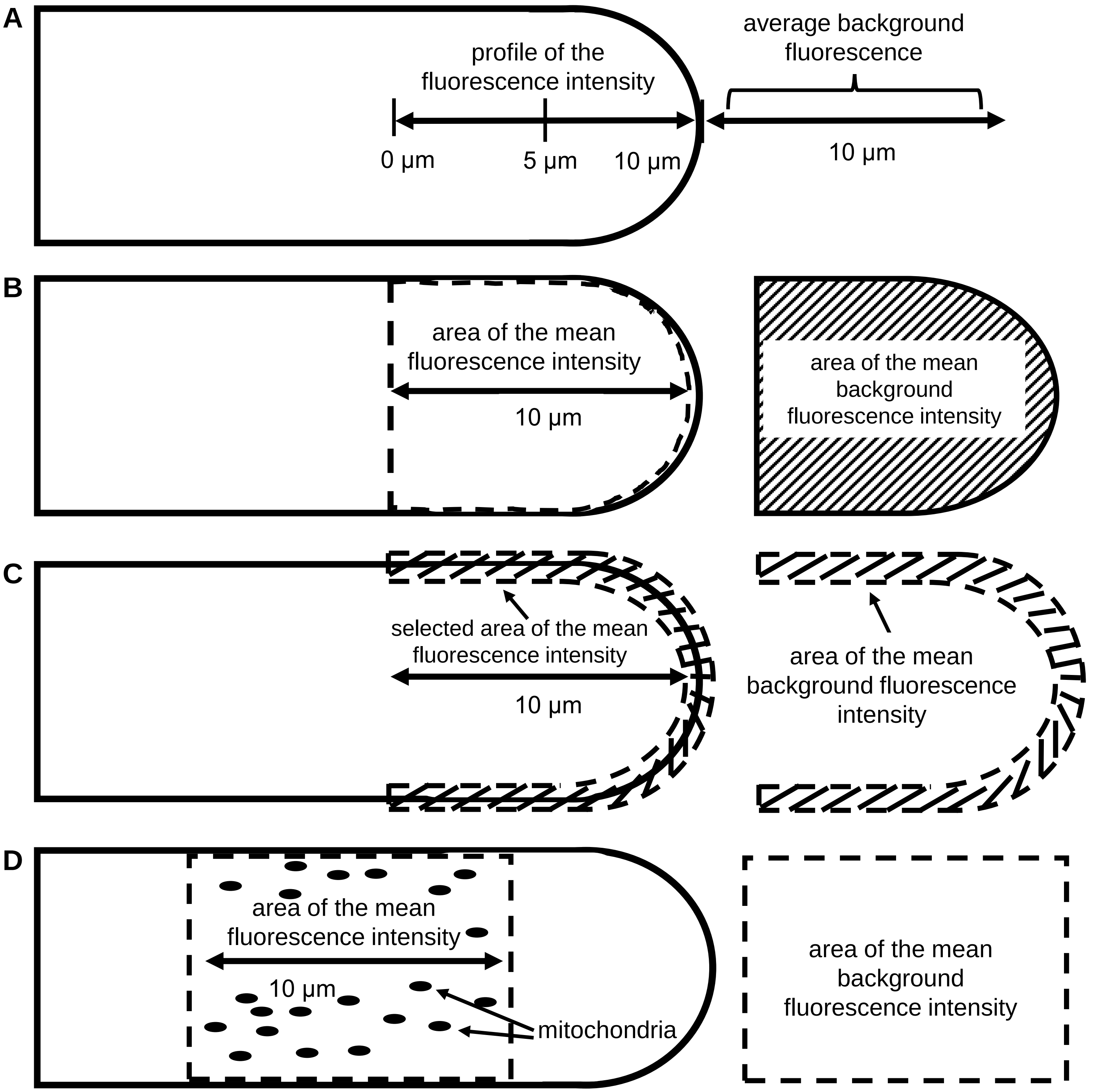


**Supplemental Figure S1. Topology of the quantitative signal intensity measurement in bulges and growing root hairs. A.** Fluorescence intensity distribution along the profile was measured in a 10 µm segment from the center of the root hair to its tip at the start (0 min), in the middle (5 min), and at the end of the scanning (10 min). **B.** The mean fluorescence intensity in area of the tip was measured in the middle of the scanning (5 min). **C.** For the Amplex$^{TM}$ Red Reagent probe, an area including only the cell wall was measured. **D.** Distribution of the CellROX$^{TM}$ Deep Red before and after valinomycin treatment was measured in area located in the subapical region of the root hair. Measurement of the background fluorescence intensity in each approach is indicated.

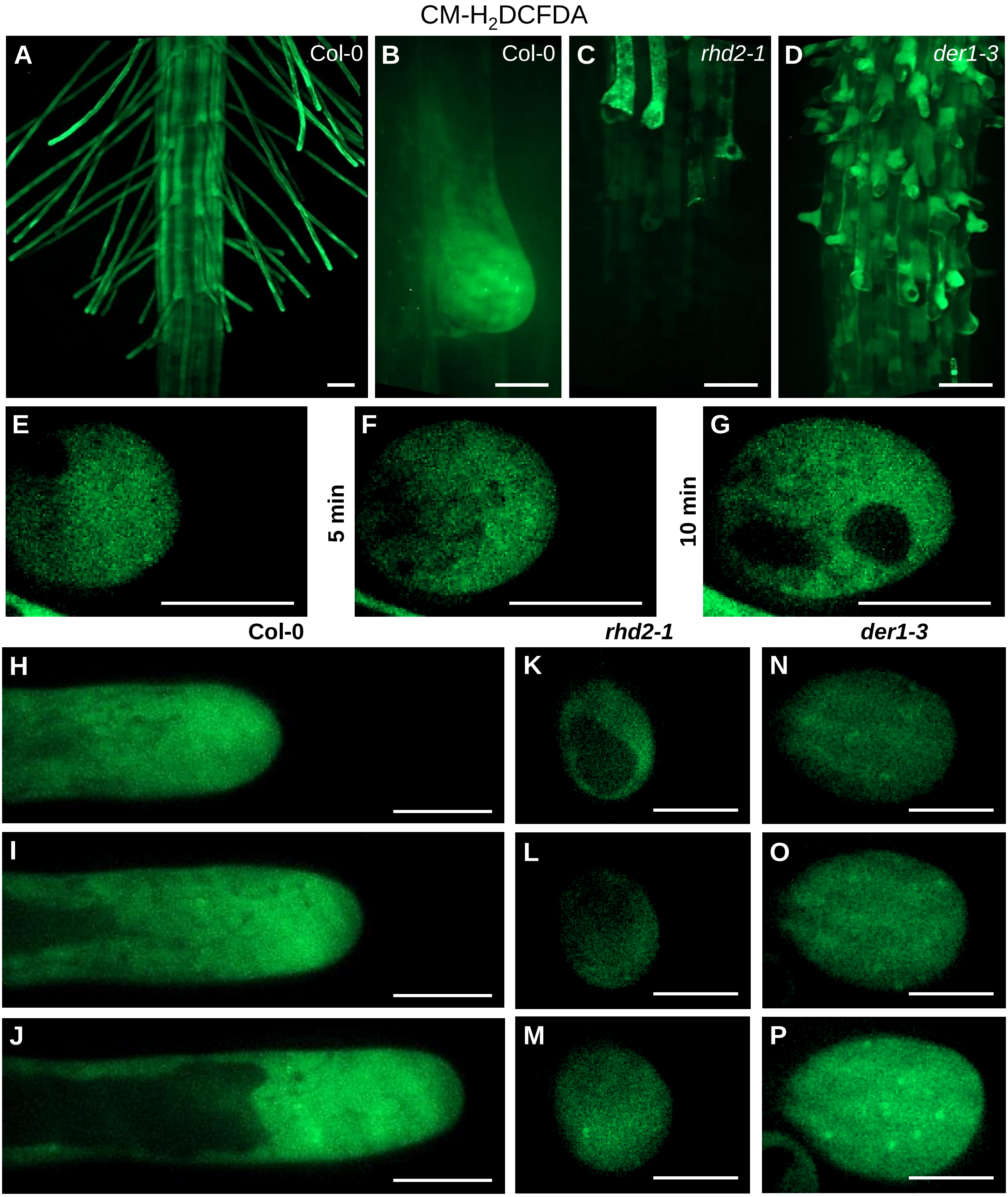


**Supplemental Figure S2. Distribution of ROS in Col-0 wild-type, *rhd2-1* and *der1-3* mutants after staining with CM-$H_2$DCFDA. A-D.** Overview of the root hair formation zone **(A)** and root hair bulge **(B)** in the root of Col-0, and the root hair formation zone with root hair bulges in roots of *rhd2-1* **(C)** and *der1-3* **(D)** mutants. **E-G.** Root hair bulges of Col-0. **H-J.** Growing root hairs of Col-0. **K-P.** Root hair bulges of *rhd2-1* **(K-M)** and *der1-3* **(N-P)** mutants. Bulges and root hairs are illustrated at time points of 0 min **(E,H,K,N)**, 5 min **(F,I,L,O)** and 10 min **(G,J,M,P)** of imaging. Image acquisition at the time point 0 min started 110 min in Col-0 bulges **(E-G)**, 154 min in Col-0 root hairs **(H-J)**, 101 min in in *rhd2-1* **(K-M)** and 49 min in *der1-3* **(N-P)** after probe application, respectively. Scale bar = 50 μm **(A,C,D)**, 10 μm **(B,E-P)**.

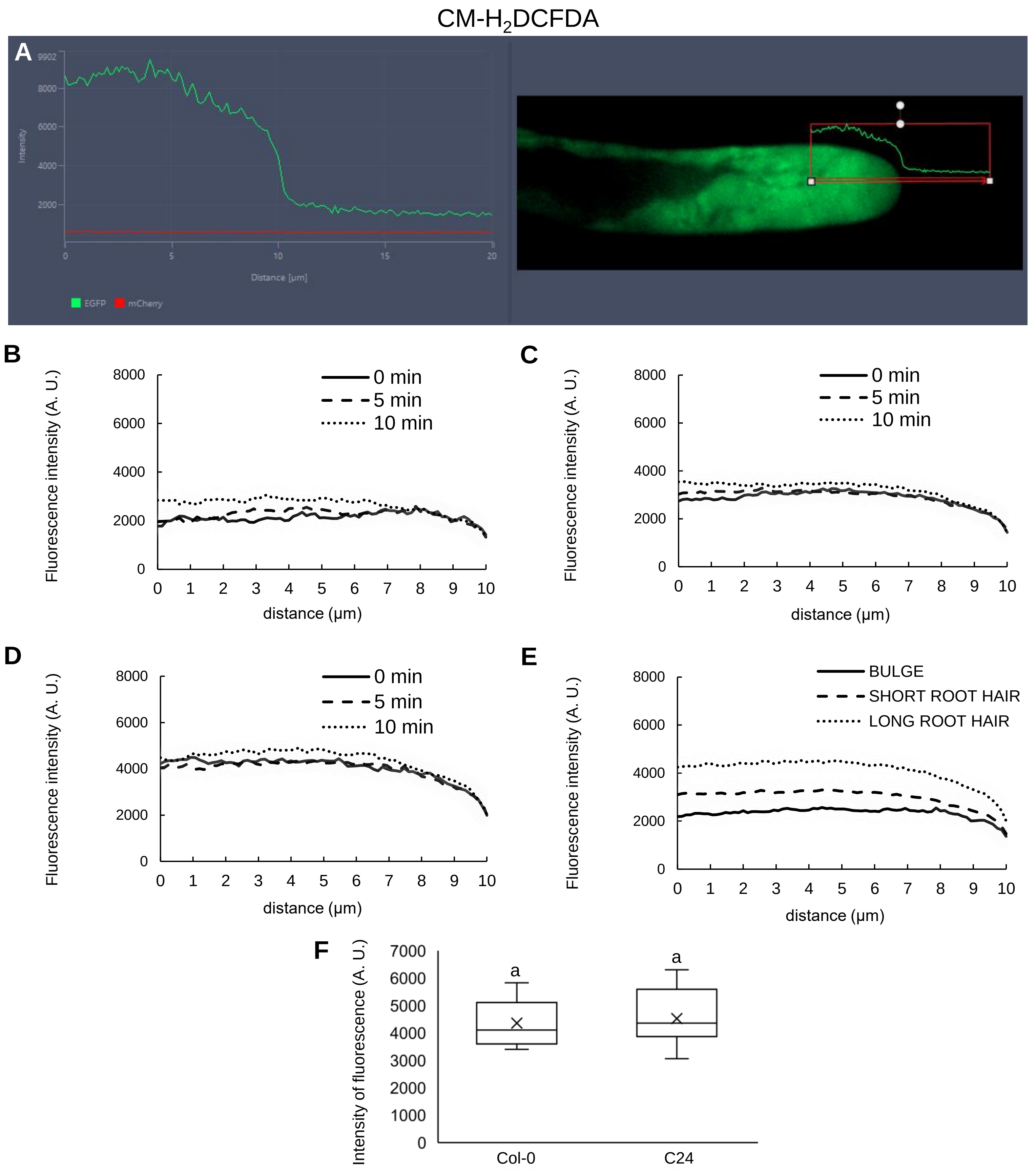


**Supplemental Figure S3. Fluorescence intensity measurements in apical parts of bulges and root hairs after ROS staining with CM-H$_2$DCFDA. A.** Presentation of fluorescence intensity measurement along a 10 µm line oriented longitudinally at the central part, reaching the apical cell wall in growing root hair (schematically illustrated in Supplemental Figure S1A) of Col-0. **B-E.** Line profile measurement of mean fluorescence intensity in bulges **(B)**, „short" **(C)** and „long" **(D)** growing root hairs. Mean fluorescence intensities were measured at 0 min, 5 min and 10 min of imaging **(B-E)**. Differences in mean fluorescence intensities of CM-H$_2$DCFDA measured during root hair development from bulges to „long" root hairs **(E)**. Growing root hairs were annotated as „short", reaching the range of 10-200 µm, and as „long", reaching the length of 200 µm and more at the time of imaging. **F.** Semi-quantitative mean CM-H$_2$DCFDA fluorescence intensity analysis (measured area schematically illustrated in Supplemental Figure S1B) in growing root hairs of Col-0 and C24 wild-types. N = 10-21. Box plots display the first and third quartiles, split by the median; the crosses indicate the mean values; whiskers extend to include the max/min values. Lowercase letters indicate statistical significance between lines according to one-way ANOVA with Fisher's LSD tests ($P < 0.05$).

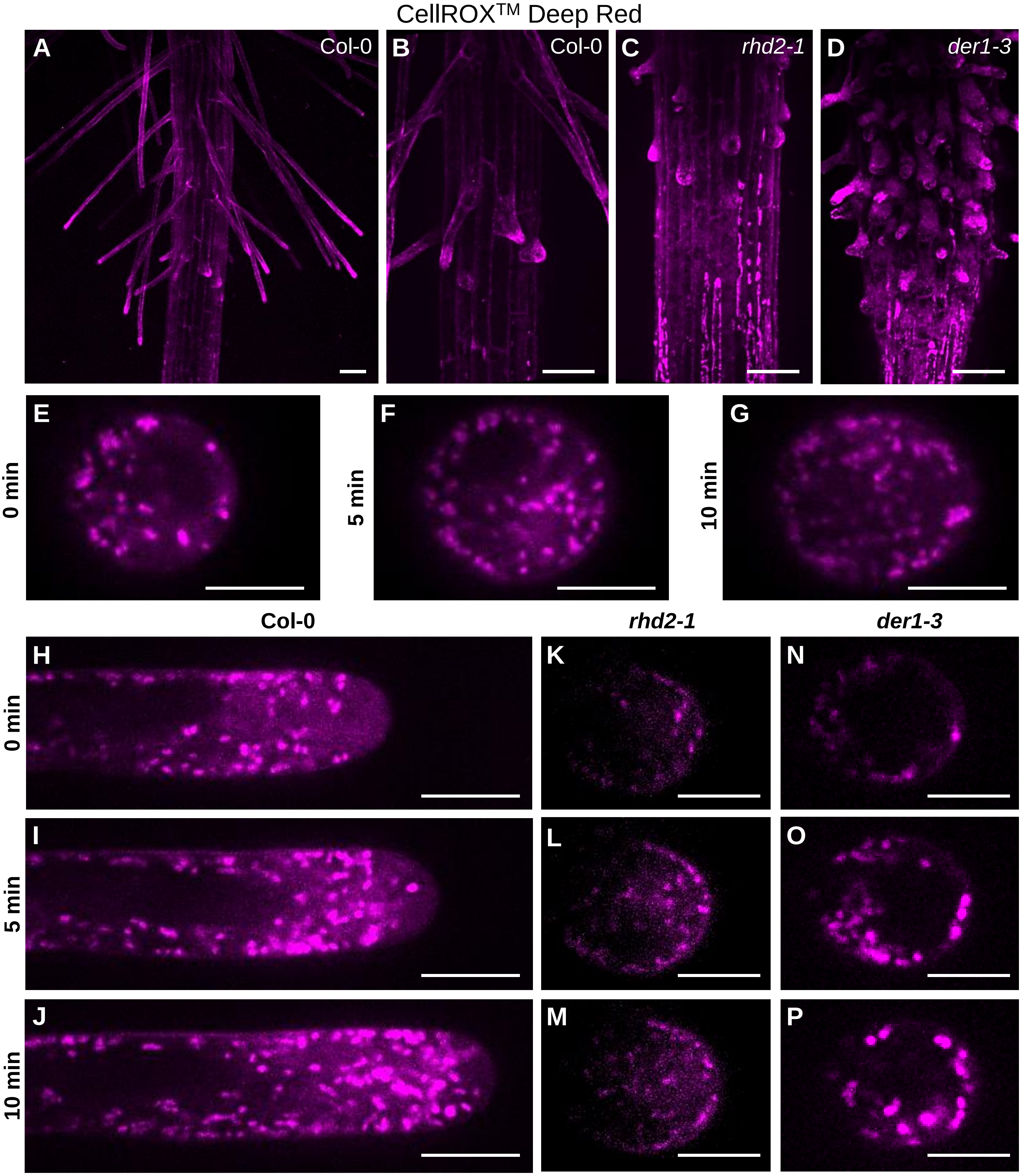


**Supplemental Figure S4. Distribution of ROS in Col-0 wild-type, *rhd2-1* and *der1-3* mutants after staining with CellROX™ Deep Red. A-D.** Overview of the root hair formation zone **(A)** and root hair bulges **(B)** in the root of Col-0, and the root hair formation zone with root hair bulges in roots of *rhd2-1* **(C)** and *der1-3* **(D)** mutants. **E-G.** Root hair bulges of Col-0. **H-J.** Growing root hairs of Col-0. **K-P.** Root hair bulges of *rhd2-1* **(K-M)** and *der1-3* **(N-P)** mutants. Bulges and root hairs are illustrated at time points of 0 min **(E,H,K,N)**, 5 min **(F,I,L,O)** and 10 min **(G,J,M,P)** of imaging. Image acquisition at the time point 0 min started 37 min in Col-0 bulges **(E-G)**, 47 min in Col-0 root hairs **(H-J)**, 74 min in in *rhd2-1* **(K-M)** and 7 min in *der1-3* **(N-P)** after probe application, respectively. Scale bar = 50 µm **(A-D)**, 10 µm **(E-P)**.

CellROX$^{TM}$ Deep Red

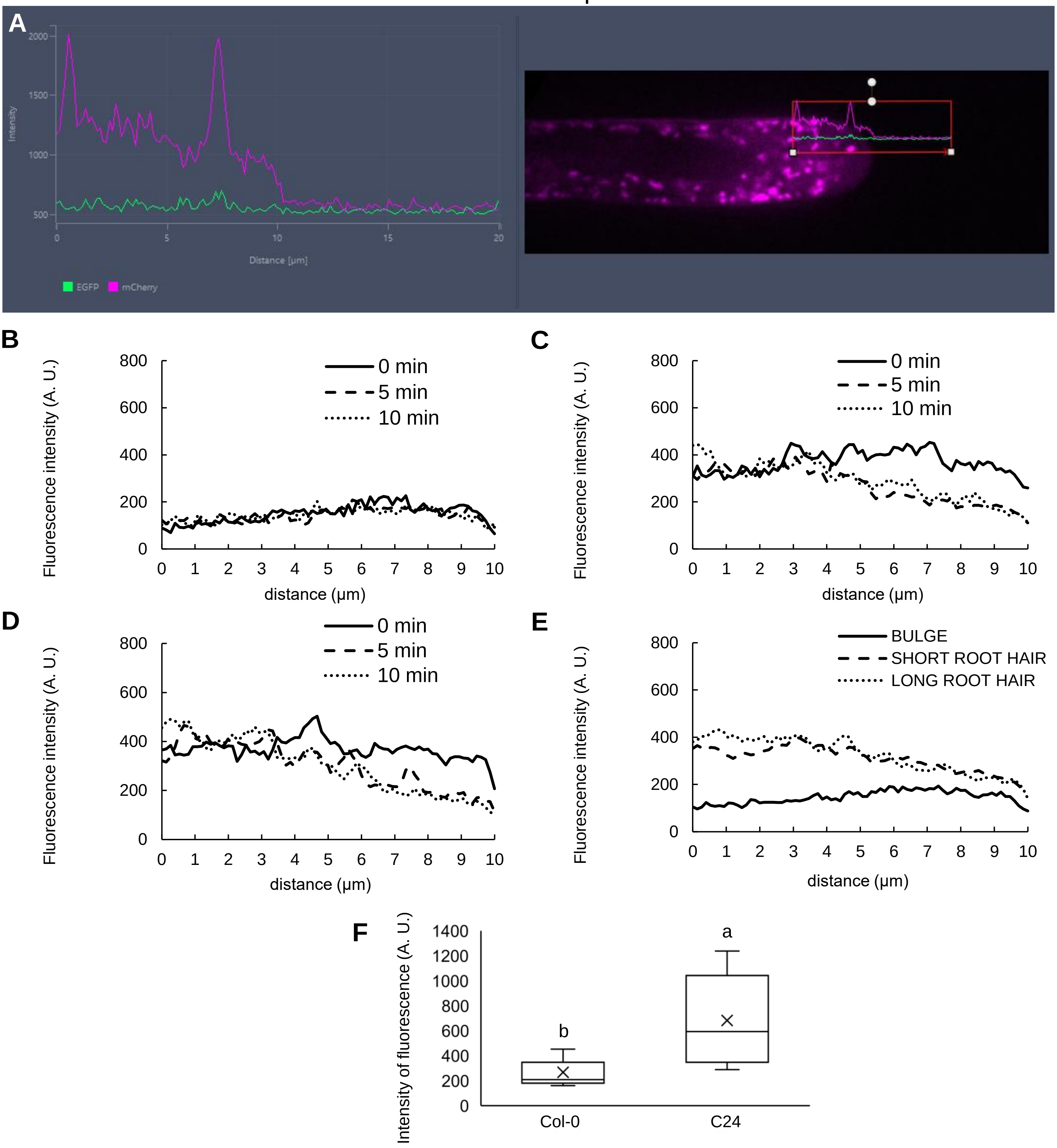


**Supplemental Figure S5. Fluorescence intensity measurements in apical parts of bulges and root hairs after ROS staining with CellROX$^{TM}$ Deep Red. A.** Presentation of fluorescence intensity measurement along a 10 µm line oriented longitudinally at the central part, reaching the apical cell wall in growing root hair (schematically illustrated in Supplemental Figure S1A) of Col-0. **B-E.** Line profile measurement of mean fluorescence intensity in bulges **(B)**, „short“ **(C)** and „long“ **(D)** growing root hairs. Mean fluorescence intensities were measured at 0 min, 5 min and 10 min of imaging **(B-E)**. Differences in mean fluorescence intensities of CellROX$^{TM}$ Deep Red measured during root hair development from bulges to „long“ root hairs **(E)**. Growing root hairs were annotated as „short“, reaching the range of 10-200 µm, and as „long“, reaching the length of 200 µm and more at the time of imaging. **F.** Semi-quantitative mean CellROX$^{TM}$ Deep Red fluorescence intensity analysis (measured area schematically illustrated in Supplemental Figure S1B) in growing root hairs of Col-0 and C24 wild-types. N = 10-17. Box plots display the first and third quartiles, split by the median; the crosses indicate the mean values; whiskers extend to include the max/min values. Lowercase letters indicate statistical significance between lines according to one-way ANOVA with Fisher’s LSD tests ($P < 0.05$).

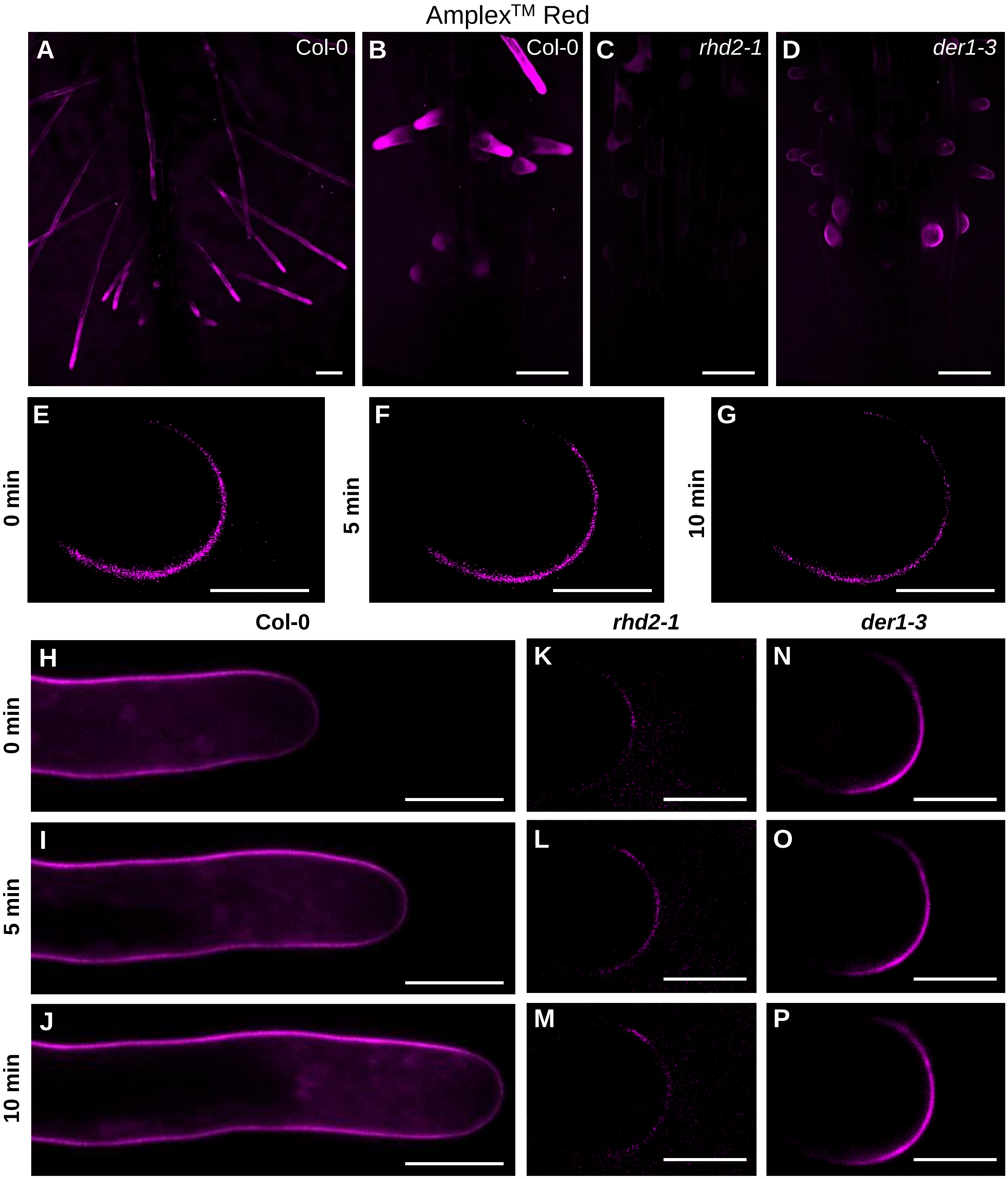


**Supplemental Figure S6. Distribution of ROS in Col-0 wild-type, *rhd2-1* and *der1-3* mutants after staining with Amplex™ Red. A-D.** Overview of the root hair formation zone **(A)** and root hair bulges **(B)** in the root of Col-0, and the root hair formation zone with root hair bulges in roots of *rhd2-1* **(C)** and *der1-3* **(D)** mutants. **E-G.** Root hair bulges of Col-0. **H-J.** Growing root hairs of Col-0. **K-P.** Root hair bulges of *rhd2-1* **(K-M)** and *der1-3* **(N-P)** mutants. Bulges and root hairs are illustrated at time points of 0 min **(E,H,K,N)**, 5 min **(F,I,L,O)** and 10 min **(G,J,M,P)** of imaging. Image acquisition at the time point 0 min started 28 min in Col-0 bulges **(E-G)**, 47 min in Col-0 root hairs **(H-J)**, 5 min in in *rhd2-1* **(K-M)** and 29 min in *der1-3* **(N-P)** after probe application, respectively. Scale bar = 50 µm **(A-D)**, 10 µm **(E-P)**.

Amplex™ Red

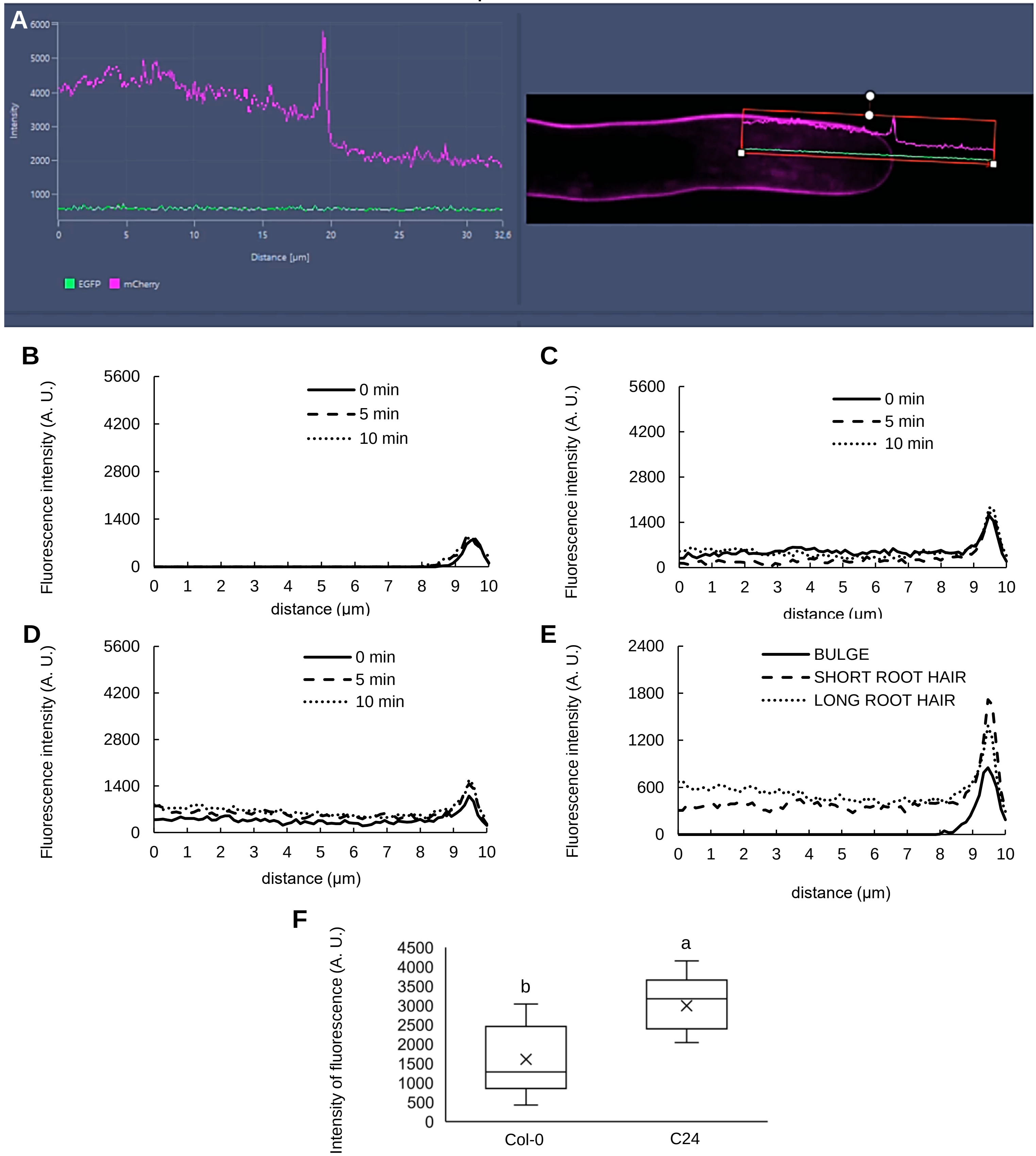


**Supplemental Figure S7. Fluorescence intensity measurements in apical parts of bulges and root hairs after ROS staining with Amplex™ Red. A.** Presentation of fluorescence intensity measurement along a 10 µm line oriented longitudinally at the central part, reaching the apical cell wall in growing root hair (schematically illustrated in Supplemental Figure S1A) of Col-0. **B-E.** Line profile measurement of mean fluorescence intensity in bulges **(B)**, „short“ **(C)** and „long“ **(D)** growing root hairs. Mean fluorescence intensities were measured at 0 min, 5 min and 10 min of imaging **(B-E)**. Differences in mean fluorescence intensities of Amplex™ Red measured during root hair development from bulges to „long“ root hairs **(E)**. Growing root hairs were annotated as „short“, reaching the range of 10-200 µm, and as „long“, reaching the length of 200 µm and more at the time of imaging. **F.** Semi-quantitative mean Amplex™ Red fluorescence intensity analysis (measured area schematically illustrated in Supplemental Figure S1B) in growing root hairs of Col-0 and C24 wild-types. N = 10. Box plots display the first and third quartiles, split by the median; the crosses indicate the mean values; whiskers extend to include the max/min values. Lowercase letters indicate statistical significance between lines according to one-way ANOVA with Fisher’s LSD tests ($P < 0.05$).

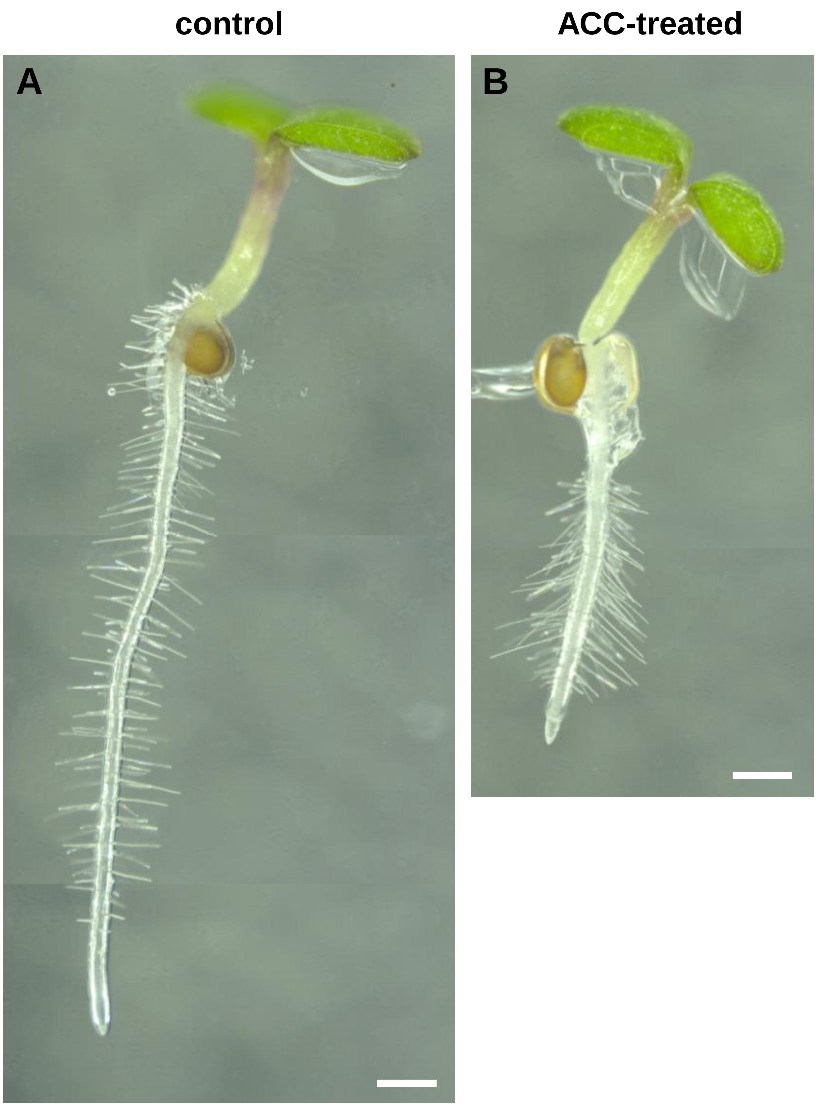


**Supplemental Figure S8. Phenotype of Col-0 wild-type seedlings treated with the ethylene precursor ACC. A-B.** Two-day-old seedlings germinating and growing on solidified control medium were transferred to solidified control medium **(A)**, or medium containing 0.7 μmol·$L^{-1}$ ACC for 24h **(B).** A composite image of three and two consequential frames is shown in **(A)** and **(B)**, respectively. Scale bar = 500 μm **(A-B)**.

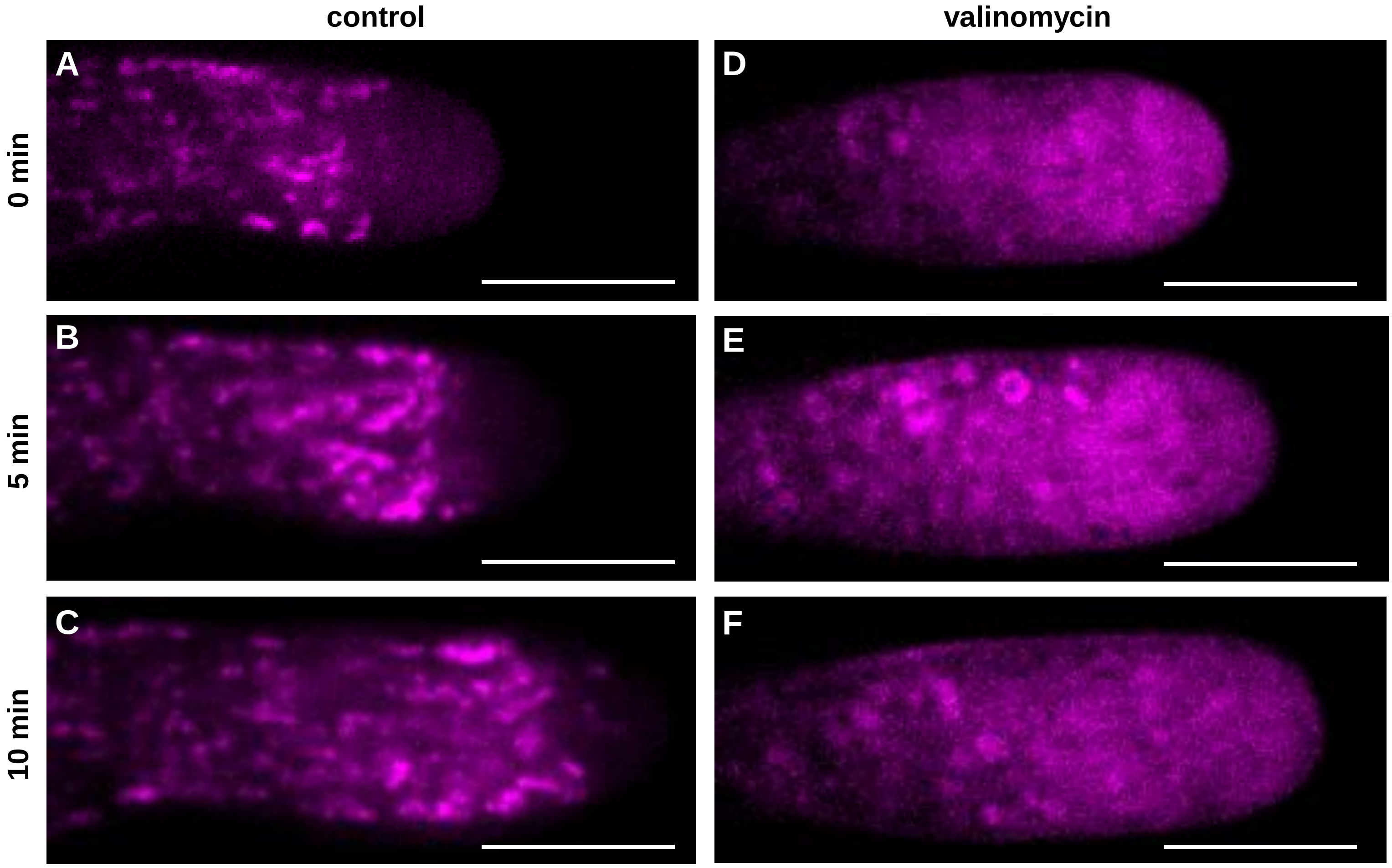


**Supplemental Figure S9. Distribution of ROS stained with CellROX$^{TM}$ Deep Red in root hairs of Col-0 after pre-treatment with the mitochondrial ionophore valinomycin. A-F.** Growing control root hair of Col-0 stained with CellROX$^{TM}$ Deep Red **(A-C)**, and root hair stained with CellROX$^{TM}$ Deep Red that was pre-treated with valinomycin **(D-F)**. Root hairs were imaged at time points of 0 min **(A,D)**, 5 min **(B,E)** and 10 min **(C,F)** of growth. Scale bar = 10 μm **(A-F)**.

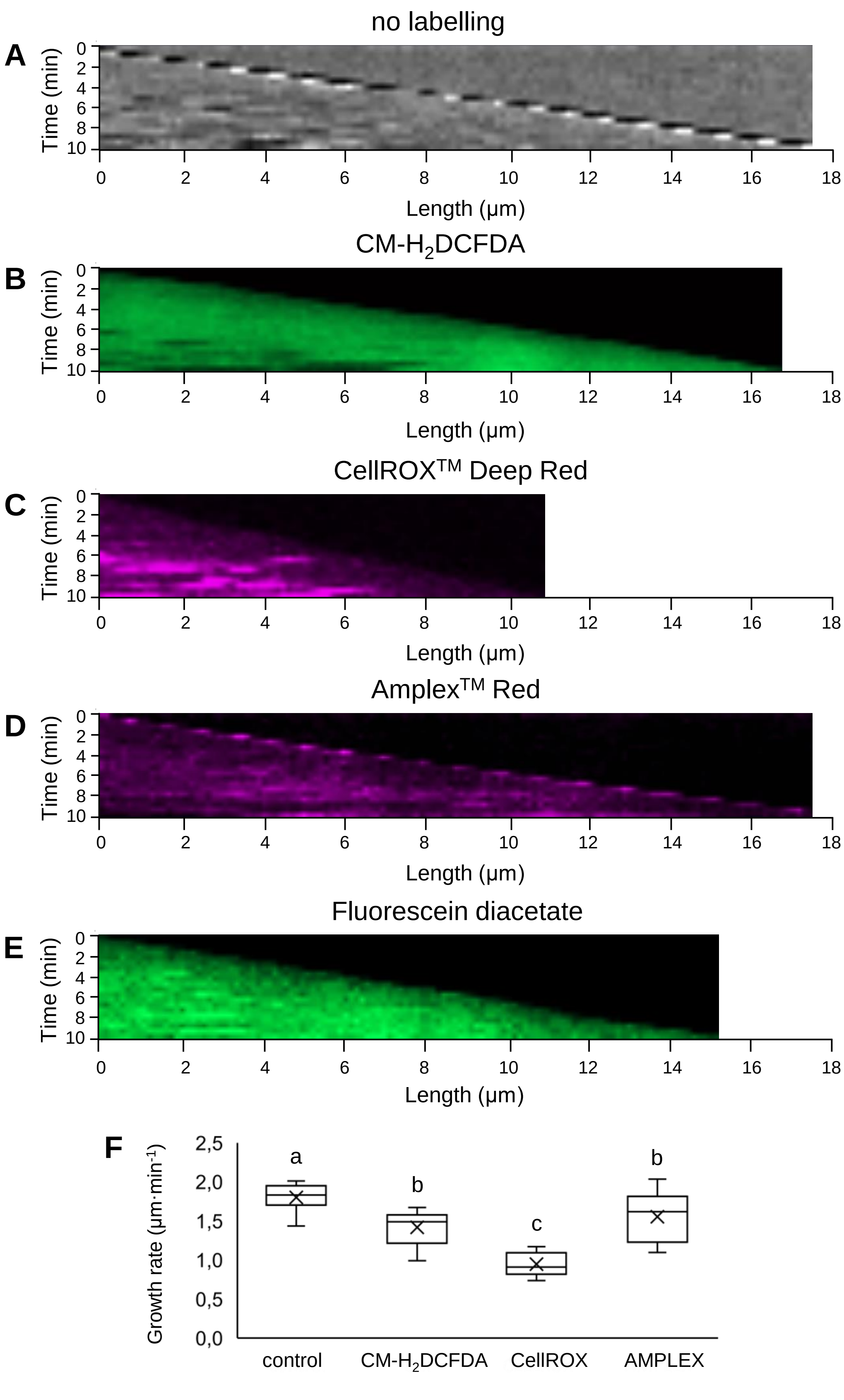


**Supplemental Figure S10. Speed of the root hair tip growth analyzed by kymographs. A-E.** Representative kymographs showing a velocity of root hair tip growth of control non-stained root hair **(A)**, CM-$H_2$DCFDA-stained root hair **(B)**, CellROX$^{TM}$ Deep Red-stained root hair **(C)**, Amplex$^{TM}$ Red-stained root hair **(D)**, and FDA-stained root hair **(E)**. **F.** Averaged root hair tip growth rate of control non-stained root hairs and root hairs stained with CM-$H_2$DCFDA, CellROX$^{TM}$ Deep Red and Amplex$^{TM}$ Red. Root hair tip growth was measured during imaging within the time period of 10 min. N = 10-21. Box plots display the first and third quartiles, split by the median; the crosses indicate the mean values; whiskers extend to include the max/min values. Lowercase letters indicate statistical significance between lines according to one-way ANOVA with Fisher's LSD tests ($P < 0.05$).
